# GESTATIONAL PROTEIN RESTRICTION PROGRAMS SEXUALLY DIVERGENT BEHAVIORAL AGING IN MICE: EARLY AFFECTIVE VULNERABILITY IN MALES AND LATE-LIFE MNEMONIC PRESERVATION AT A HEDONIC COST IN FEMALES

**DOI:** 10.64898/2026.11.17.765387

**Authors:** Vinicius Schiavinatto Mariano, Amanda Cristina de Souza, Gabriel Boer Grigoletti-Lima, José Antonio Rocha Gontijo, Patrícia Aline Boer

## Abstract

Maternal protein restriction is a well-established model of developmental programming, yet its consequences for behavior and cognition across the second half of life, and how these differ between sexes, remain poorly characterized. We fed C57BL/6J dams a normal-protein (NP, 17%) or low-protein (LP, 6%) diet throughout gestation and followed male and female offspring with an integrated battery assessing reward sensitivity, locomotion, anxiety-like behavior, social behavior, recognition memory, spatial learning and cognitive flexibility at 12 (middle age) and 18 months (old age). Maternal care was quantified from postnatal days 2 to 9. LP offspring of both sexes were born lighter and caught up by postnatal day 7. LP dams spent more time self-grooming, with longer self-grooming bouts, and took longer to retrieve their pups, whereas licking/grooming, its fragmentation and nursing were unchanged. Middle-aged LP males showed anhedonia-like behavior, anxiety-like behavior in the elevated plus-maze, reduced exploration and impaired short-term object recognition; in old age, affective indices normalized, but males became hyperactive and used less efficient search strategies in the Barnes maze. Middle-aged LP females showed hyperactivity, stereotypy and thigmotaxis; at 18 months they developed anhedonia-like behavior yet retained long-term object, social and spatial memory that had declined in aged NP females. A qualitative comparison with a prenatal-stress cohort tested in the same laboratory with the same battery revealed an overlapping, sex- and age-specific signature, particularly for the timing of anhedonia and for late-life memory outcomes. Gestational protein restriction, therefore, programs domain-specific and sexually divergent behavioral aging trajectories rather than uniform cognitive decline. These findings support sex-stratified, life-course designs in fetal programming research.

## 1. INTRODUCTION

The prevalence of nutritional adversities during pregnancy remains high worldwide. The “Developmental Origins of Health and Disease” (DOHaD) theory, derived from epidemiological data, posits that conditions experienced in utero shape susceptibility to chronic diseases throughout the lifespan (Barker, 1995; Gluckman & Hanson, 2004; Hales & Barker, 1992). Among experimental models, maternal protein restriction in rodents has been a specific focus. Our group has characterized a model using a 6% protein diet during gestation, in which we observed reduced birth weight and a lower number of nephrons, followed by the development of hypertension in the offspring (Lamana et al., 2021; Sene et al., 2021). More recently, using this same model, we observed an acceleration of aging-related phenotypes in peripheral organs, such as bones and kidneys (Calsa et al., 2026; Pires et al., 2026). Regarding neural consequences, we found that gestational protein restriction led to dendritic atrophy in the hippocampal CA3 region—without detectable changes in learning and memory performance in adult male rats—demonstrating that a structural modification does not, by itself, determine a deficit in a given task (Lopes et al., 2013). Conversely, our studies in aged males identified alterations in spatial navigation and hippocampal proteins, alongside the preservation of certain measures of working memory (Grigoletti-Lima et al., 2022). However, it remains unclear whether gestational protein restriction shapes the trajectory of brain aging in a sex-specific manner. Various mechanisms can link maternal protein restriction to neurodevelopment. In rats, maternal protein restriction reduces the expression of placental 11β-HSD2 and promotes a persistent increase in the expression of glucocorticoid receptors (GR) across several tissues—both fetal and adult—including the brain (Bertram et al., 2001; Togher et al., 2014). Using the same model employed here, our group reported a reduction in the levels of glucocorticoid receptors (GR) and mineralocorticoid receptors (MR), as well as alterations in monoamine levels, within the amygdala and the bed nucleus of the stria terminalis (BNST) during the development of male rats (Torres et al., 2023, 2024). Furthermore, in adult males, we observed increased basal corticosterone, reduced GR levels in the hippocampus, and an increase in anxiety-like behaviors (Mariano et al., 2025). Early-life protein restriction has also been associated with anhedonia-like behaviors linked to reduced BDNF (brain-derived neurotrophic factor) signaling via the TrkB receptor in the hippocampus (Gutiérrez et al., 2025), alterations in midbrain dopaminergic circuits (Prodani et al., 2024), and sex-specific hippocampal astrogliosis (Bello-Medina et al., 2026).

The postnatal environment constitutes a second, often overlooked, pathway for programming. Maternal behaviors, such as nursing in an arched-back posture and licking/grooming offspring, increase the expression of glucocorticoid receptors (GR) in the hippocampus and enhance negative feedback within the hypothalamic-pituitary-adrenal (HPA) axis in the offspring (Buschdorf & Meaney, 2016; Caldji et al., 2011). Conversely, fragmented and unpredictable maternal care is associated with adverse emotional outcomes later in life (Baram et al., 2012; Glynn & Baram, 2019; Molet et al., 2016). Gianatiempo et al. (2020) observed that female mice subjected to protein restriction provided less care to their offspring, with such care being more fragmented. Furthermore, Alberca et al. (2024) noted anxiety-like and anhedonic-like behaviors in female mice—though, in our own model involving dams subjected to gestational protein restriction, we did not detect any alterations in maternal care (Mariano et al., 2025). Given its role in modulating developmental programming and the existing discrepancy in findings, characterizing maternal behavior is essential for interpreting the phenotypes observed in the offspring. Most behavioral studies on early-life protein restriction have focused on young adults, examined only one sex, or involved exposure periods that spanned both gestation and lactation (Belluscio et al., 2014; Crossland et al., 2017; Fesser et al., 2021). The few available studies that track the life cycle—combining dietary restriction during gestation and lactation—describe persistently low levels of exploration, anxiety-like behaviors, and impairments in working memory during middle age, as well as premature aging of spatial memory in females (Ferroni et al., 2022, 2023). The sex-dependent effects of prenatal malnutrition remain a subject of debate: a systematic review found little consistent evidence for greater vulnerability in males and noted that fewer than half of the studies formally tested for interactions between sex and diet (Christians et al., 2021). In humans, exposure to famine during early gestation has been associated with poorer performance in selective attention at 56–59 years of age, suggesting accelerated cognitive aging (de Rooij et al., 2010).

Three gaps, therefore, motivated the present study: first, the scarcity of data on behavioral outcomes resulting from protein restriction limited to the gestational period, assessed in middle and old age; second, the lack of experimental designs that analyze both sexes throughout these life stages within a single model; and third, the absence of integrated phenotyping that combines maternal care with domains such as reward, anxiety, social behavior, recognition, and spatial memory in the offspring. We characterized the offspring (males and females) of the C57BL/6J strain, born to dams fed either a normoprotein (NP) or a low-protein (LP) diet, at 12 and 18 months of age, using a battery of tests identical to that recently applied by our group in a model of unpredictable chronic mild stress (UCMS) during the prenatal period (de Souza et al., 2026, submitted; doi.org/10.64898/2026.10.09.757621). This parallel design allowed for the exploration of an additional question: do nutritional and psychological adversities during the prenatal period converge toward a common behavioral signature of aging? Our hypothesis was that gestational protein restriction would produce effects that are both sex-specific and age-dependent.

## 2. MATERIALS AND METHODS

### 2.1. Ethical statement and animal housing

All experimental procedures complied with the guidelines of the Brazilian National Council for the Control of Animal Experimentation (CONCEA) and were approved by the Ethics Committee for Animal Research of the State University of Campinas (CEUA 5776-1/2021). Mice were housed in standard polypropylene cages (40 × 20 × 15 cm) with wood-shaving bedding under controlled conditions (21 ± 2 °C; 12:12 h light–dark cycle, lights on at 06:00 h) with *ad libitum* access to water and chow (Nuvital®, Colombo, PR, Brazil). The overall experimental design is summarized in Supplementary Fig. S1.

### 2.2. Animals, breeding and diets

Male (n = 20) and female (n = 40) C57BL/6J mice were obtained from the Multidisciplinary Center for Biological Research (CEMIB, UNICAMP). At six weeks of age, females were paired with males (2:1) in breeding cages. Pregnancy was confirmed by the presence of a vaginal plug, designated gestational day 0 (GD0); males were then removed and females were housed individually. Pregnant dams received either a normal-protein diet containing 17% protein (NP, control) or a low-protein diet containing 6% protein (LP) throughout gestation.

Both diets were isocaloric, casein-based modifications of the AIN-93G formulation (PragSoluções Biociências, SP, Brazil; Supplementary Table S1). Fifty-two females were allocated to the NP diet and 53 to the LP diet; pregnancy was confirmed in 20 NP and 30 LP dams. From parturition to weaning, all dams received standard laboratory chow (Nuvital^®^). Weekly dam’s weight and intake were measured. Litter size was 7.1 ± 0.8 (NP) and 6.5 ± 0.7 (LP) pups. Perinatal losses included total cannibalism (NP 1; LP 5), stillbirth or death of the entire litter (NP 2; LP 3) and partial cannibalism (NP 1; LP 5), so that 12 NP and 10 LP litters were used in the study.

### 2.3. Offspring biometric parameters

Offspring of both sexes were weighed at birth (postnatal day 0, PND0) and at PND7, PND14 and PND21 to track postnatal growth. At PND1, litters were culled to a maximum of eight pups to standardize access to milk.

### 2.4. Maternal behavior

Maternal behavior was assessed following Molet et al. (2016) and Zoubovsky et al. (2022) from PND2 to PND9, in two daily 30-min sessions during the light phase (09:00–10:30 and 14:00–15:30), after 30 min of acclimation to the observation room and without cage changes. The duration (s) of the following behaviors was recorded: licking/grooming (LG), arched-back nursing (ABN), passive nursing (PN), time off the nest (ON), nest building (NB), self-grooming (SG) and eating/drinking. Four independent observers recorded behaviors with the smartphone application Countee (Slanzi & Fernand, 2024), and inter-observer agreement was checked with the intraclass correlation coefficient on a subset of recordings. Maternal behavior was analyzed in 8 NP and 10 LP dams, and pup retrieval in 12 NP and 13 LP dams. Outliers were identified with the ROUT method (Q = 1%) and removed for individual parameters, which accounts for the variation in sample size across panels in Fig. 2.

#### Behavioral fragmentation

Fragmentation of maternal care was estimated as a marker of unpredictability (Baram et al., 2012; Molet et al., 2016) from the number of bouts and the mean bout duration (s) of licking/grooming and self-grooming, with shorter and more frequent bouts denoting more fragmented behavior.

#### Pup retrieval test

At PND3, adapted from Wang and Storm (2011), the dam was briefly removed and three randomly selected pups were placed on the side of the cage opposite to the nest (two in the corners and one in the middle). The dam was returned between the nest and the pups, and the latency to retrieve the first pup and all three pups to the nest was recorded (maximum 600 s).

### 2.5. Behavioral testing of the offspring

After weaning at PND21, offspring were housed in same-sex groups of three to five per cage until testing at 12 months (middle age) and 18 months (old age). One offspring per dam and sex was assigned to each experimental group, so that the dam (litter) was the experimental unit. To limit carry-over effects, animals were allocated to two independent cohorts: Cohort 1 completed only the Barnes maze protocol (18 months), and Cohort 2 underwent a sequential battery comprising the sucrose preference test (SPT), open field, novel object recognition (NOR), elevated plus-maze (EPM) and three-chamber social test. Testing was performed during the light phase (08:00–14:00) in a dedicated room (22 ± 1 °C; background noise < 50 dB; 30–50 lux unless otherwise specified), after ≥ 30 min of acclimation. Apparatuses were cleaned with 70% ethanol between trials, and testers were blinded to group allocation whenever possible. Group sizes for each test are given in Table 1.

**Table 1.** Experimental groups, group sizes (NP/LP) and statistical tests for each outcome. Group sizes as reported in the official Methods/Results. Dam numbers for maternal behavior varied by parameter (Fig. 2). In the official Methods, offspring were assigned one per dam to each group; the dam was therefore the experimental unit.

| **Outcome** | **Females 12 mo** | **Females 18 mo** | **Males 12 mo** | **Males 18 mo** | **Analysis** |
|---|---|---|---|---|---|
| Offspring body weight, PND0–21 (Fig. 1) | 35/27 (all females) | — | 29/35 (all males) | — | Two-way ANOVA |
| Maternal behavior, PND2–9 (Fig. 2A–J) | Dams: 8/8-10 |  |  |  | Student's t-test |
| Pup retrieval, PND3 (Fig. 2K–L) | Dams: 9/11-13 |  |  |  | Student's t-test |
| Sucrose preference (Fig. 4A, E) | 17/17 | 14/16 | 10/11 | 15/12 | Multiple t-tests (Holm–Šidák) |
| Open field (Fig. 3) | 9–13/8–9 | 12–14/11–15 | 10–11/10–11 | 12–13/9–12 | Multiple t-tests |
| Novel object recognition (Figs. 6–7) | 14/13 | 13/15 | 12/11 | 9/11 | Two-way ANOVA; t-test (DI) |
| Elevated plus-maze (Fig. 4B–D, F–H) | 11/18 | 14/16 | 8/10 | 14/11 | Multiple t-tests |
| Three-chamber social test (Fig. 5) | 13/12 | 15/12 | 12/10 | 15/12 | Two-way ANOVA; t-test (indices) |
| Barnes maze (Figs. 8–9) | — | 14/15 | — | 14/15 | Two-way RM ANOVA; t-test / Mann–Whitney |

#### 2.5.1. Sucrose preference test

Anhedonia-like behavior was assessed in individually housed animals over three consecutive days: two water bottles on day 1 (habituation), and one water and one 1% (w/v) sucrose bottle on days 2 and 3, with bottle positions swapped on day 3. In line with current recommendations (Berrio et al., 2024), animals were not food- or water-deprived. Sucrose preference (%) = sucrose intake / (sucrose + water intake) × 100; total fluid intake was also recorded.

#### 2.5.2. Open field (activity monitor)

Locomotion and exploration were recorded for 5 min under dim light (30 lux) in an infrared activity monitor (45 × 45 × 45 cm; ENV-515, Med Associates Inc., St. Albans, VT, USA) equipped with two 16-beam arrays. Activity time, distance travelled, velocity, rearing (standing) time, stereotyped movements and time spent at the borders (thigmotaxis) and in the center were analyzed. Group sizes, which varied across parameters after outlier exclusion, are given in Table 1.

#### 2.5.3. Novel object recognition

After a 5-min habituation to the empty arena (40 × 40 × 40 cm), animals explored two identical objects for 5 min (familiarization). Short-term memory was tested 3 h later and long-term memory 24 h later by replacing one familiar object with a novel object (5 min each), with object identity and position counterbalanced (Antunes & Biala, 2012). Exploration was defined as directing the nose towards the object at ≤ 2 cm. The discrimination index (DI) was computed as (time exploring the novel object − time exploring the familiar object) / total exploration time; relative exploration times of each object were also compared within groups.

#### 2.5.4. Elevated plus-maze

Anxiety-like behavior was assessed for 5 min under dim red light (5 lux) in a maze with two open arms (30 × 5 cm) and two closed arms (30 × 5 × 15 cm) elevated 50 cm above the floor (Walf & Frye, 2007). Time spent in the open and closed arms and the number of arm entries were analyzed; total entries were used as an index of general activity.

#### 2.5.5. Three-chamber social test

After a 10-min habituation with empty wire cups, the sociability phase (5 min) offered an unfamiliar age- and sex-matched conspecific (stranger 1, S1) versus an empty cup (E); in the social novelty phase (5 min) a second unfamiliar conspecific (S2) was added to the previously empty cup. Each phase was preceded by 3 min in the closed central chamber, after a 45-s interval. The sociability index was computed as S1 / (S1 + E) and the social novelty index as S2 / (S1 + S2), based on investigation times; Fig. 5 presents both indices as percentage differences between stimuli.

#### 2.5.6. Barnes maze

Spatial learning, memory and cognitive flexibility were assessed at 18 months on a circular platform (90 cm diameter) with 20 holes (5 cm) and an escape box (9 × 20 × 9 cm) beneath the target hole, under bright light (1200 lux), with four distal visual cues. After habituation on day 0 (1 min of free exploration followed by 2 min in the escape box), animals underwent five acquisition days (days 1–5; three trials per day, 180 s maximum, 15-min inter-trial interval), a 90-s probe trial without the escape box (day 6), three reversal-learning days with the escape box relocated 180° (days 7–9) and a final 90-s reversal probe (day 10). The percentage of probe time spent in the target quadrant indexed spatial memory retention. Search trajectories were classified with the Barnes maze unbiased strategy (BUNS) classification algorithm (Illouz et al., 2016) into six strategies (direct, corrected, long correction, focused search, serial and random), and each trial was assigned a cognitive score reflecting strategy efficiency: direct = 1, corrected = 0.75, long correction and focused search = 0.5, serial = 0.25, and random = 0.

### 2.6. Statistical analysis

Data are presented as mean ± standard deviation (SD) unless otherwise indicated. Normality and homogeneity of variances were tested with the Shapiro–Wilk and Levene tests, respectively. Two-group comparisons used Student’s *t*-test (or Mann–Whitney U test for non-parametric data); multiple *t*-tests were corrected with the Holm–Šidák method, and repeated or factorial designs (growth curves, Barnes acquisition and reversal latencies, within-group stimulus preferences) were analyzed by two-way analysis of variance (ANOVA) followed by Tukey’s post hoc test. Significance was set at *p* < 0.05, and 0.05 ≤ *p* < 0.10 was described as a trend. Analyses were performed in GraphPad Prism 10 (GraphPad Software, San Diego, CA, USA). To support interpretation, we additionally (i) computed, for each comparison, the minimum effect size detectable with 80% power at a two-sided α of 0.05 given the reported group sizes (sensitivity analysis; Supplementary Fig. S2), and (ii) summarized all reported contrasts in an integrated phenotypic map (Fig. 10) and in a descriptive cross-model comparison with the prenatal UCMS cohort (Fig. 11). Neither synthesis constitutes an additional inferential test.

## 3. RESULTS

### 3.1. Dam’s weight and intake and Offspring biometric parameters

LP dam’s weight was reduced at 12GD and 0PND (Supplementary Table S2). LP offspring were significantly lighter at birth than NP offspring in both females (NP n = 35; LP n = 27; *p* < 0.0001) and males (NP n = 29; LP n = 35; *p* < 0.0001) (Fig. 1A, B), confirming intrauterine growth restriction. Body weight did not differ between groups at PND7, PND14 or PND21, indicating complete catch-up growth within the first postnatal week (Fig. 1).

**Figure 1.**
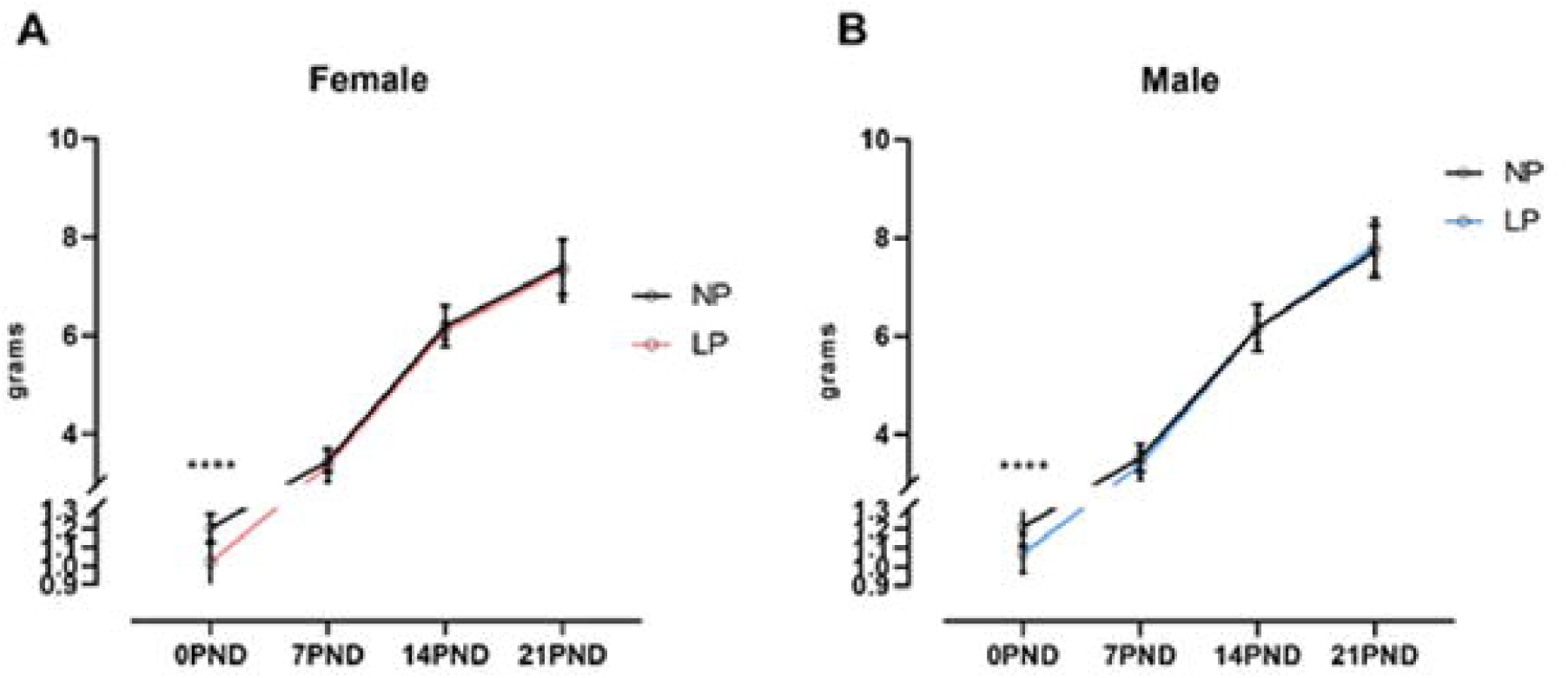
Offspring body weight from birth to weaning. (A) Female and (B) male offspring of dams fed a normal-protein (NP, 17%) or low-protein (LP, 6%) diet during gestation, weighed at postnatal days (PND) 0, 7, 14 and 21 (females: NP n = 35, LP n = 27; males: NP n = 29, LP n = 35). Data are mean ± SD. Two-way ANOVA: ****p < 0.0001 (LP vs NP at PND0).

### 3.2. Maternal behavior

#### 3.2.1. Duration and organization of maternal behaviors

LP dams spent significantly more time self-grooming than NP dams (NP 537 ± 457 s, n = 8; LP 936 ± 314 s, n = 10; *p* = 0.043; Fig. 2E), and their self-grooming bouts were longer (NP 30.6 ± 22.4 s; LP 54.7 ± 11.8 s; *p* = 0.010; Fig. 2F). In contrast, the total duration of licking/grooming (*p* = 0.27; Fig. 2A), the mean duration of licking/grooming bouts (*p* = 0.85; Fig. 2B) and the number of licking/grooming bouts (*p* = 0.61; Fig. 2C) did not differ between groups, indicating that neither the amount nor the fragmentation of licking/grooming was altered. Time off the nest (*p* = 0.70; Fig. 2D), arched-back nursing (*p* = 0.34; Fig. 2G), passive nursing (p = 0.56; Fig. 2H), eating/drinking (*p* = 0.37; Fig. 2I) and nest building (*p* = 0.27; Fig. 2J) were also unchanged.

**Figure 2.**
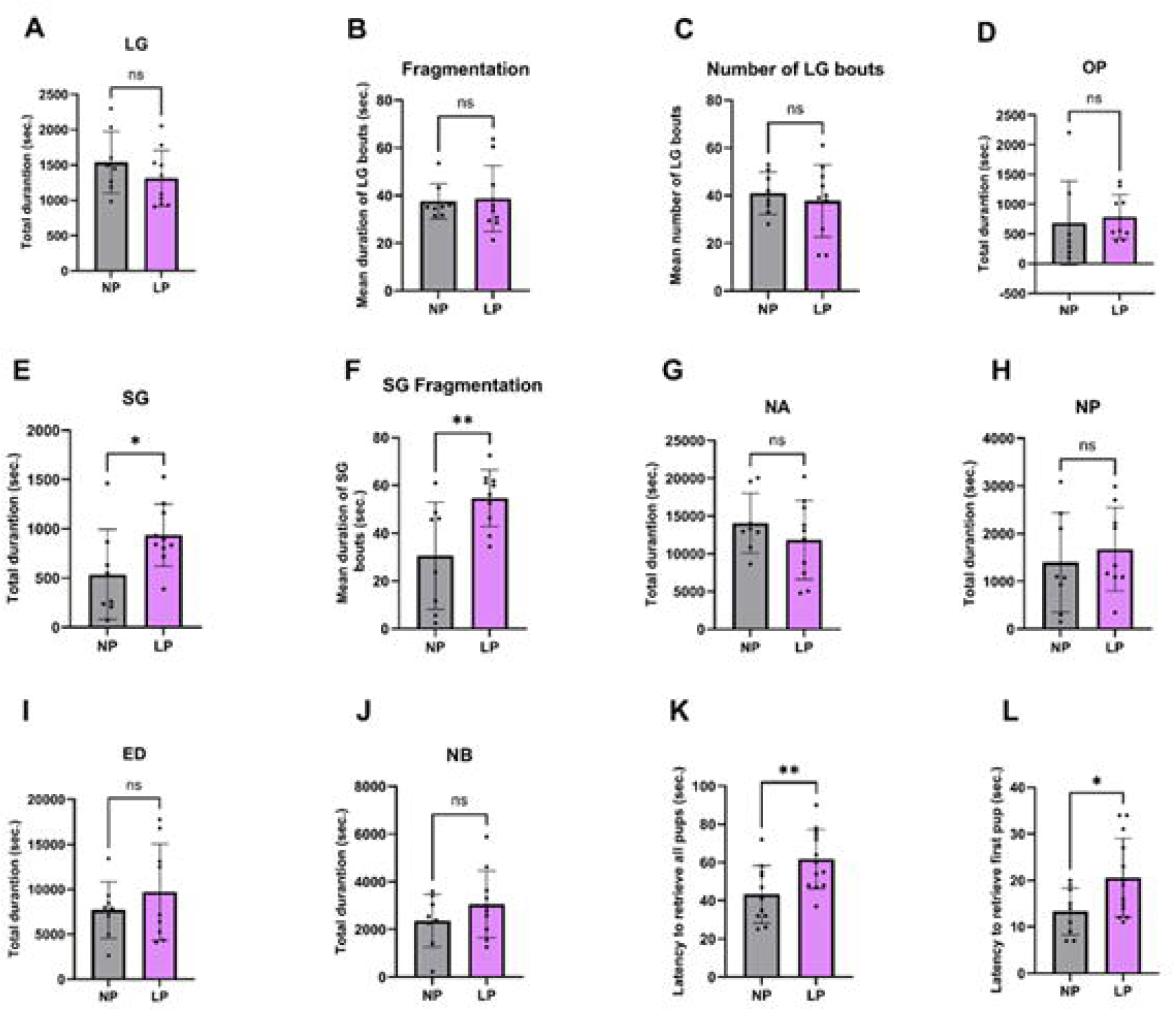
Maternal behavior from PND2 to PND9 and pup retrieval at PND3. Total duration of (A) licking/grooming; (B) mean duration of licking/grooming bouts (fragmentation); (C) number of licking/grooming bouts; total duration of (D) time off the nest, (E) self-grooming, (F) arched-back nursing, (G) passive nursing, (H) eating/drinking and (I) nest building; latency to retrieve (J) all three pups and (K) the first pup. Dots represent individual dams (n = 4–7 per group). Data are mean ± SD. Student’s t-test: *p < 0.05.

#### 3.2.2. Pup retrieval

LP dams took significantly longer than NP dams to retrieve all three pups (NP 43.3 ± 15.0 s, n = 11; LP 61.8 ± 15.4 s, n = 13; *p* = 0.007; Fig. 2K) and to retrieve the first pup (NP 13.3 ± 5.1 s, n = 9; LP 20.6 ± 8.4 s, n = 13; *p* = 0.032; Fig. 2L). Thus, gestational protein restriction increased and prolonged self-directed behavior and delayed pup-directed responses, without altering licking/grooming or nursing.

### 3.3. Locomotion and exploration in the open field

Open-field behavior depended on both sex and age (Fig. 3). At 12 months, LP females showed greater activity time, distance travelled and velocity than NP females (all *p* < 0.05; Fig. 3A–C), together with more stereotyped movements (Fig. 3E), more time at the borders and less time in the center of the arena (Fig. 3F, G), a pattern of increased thigmotaxis that is visible in the representative tracks (Fig. 3H). At 18 months, locomotor parameters and stereotypy no longer differed between groups, but LP females still spent more time at the borders and less time in the center (Fig. 3F, G). Thigmotaxis was thus the only open-field alteration that persisted across both ages in LP females. Visual inspection of Fig. 3A–C suggests that the convergence at 18 months reflected an age-related increase in activity in NP females, while LP females showed similar values at both ages; because age was not modelled as a factor, this observation remains descriptive. Rearing (standing) time did not differ between groups at either age (Fig. 3D).

**Figure 3.**
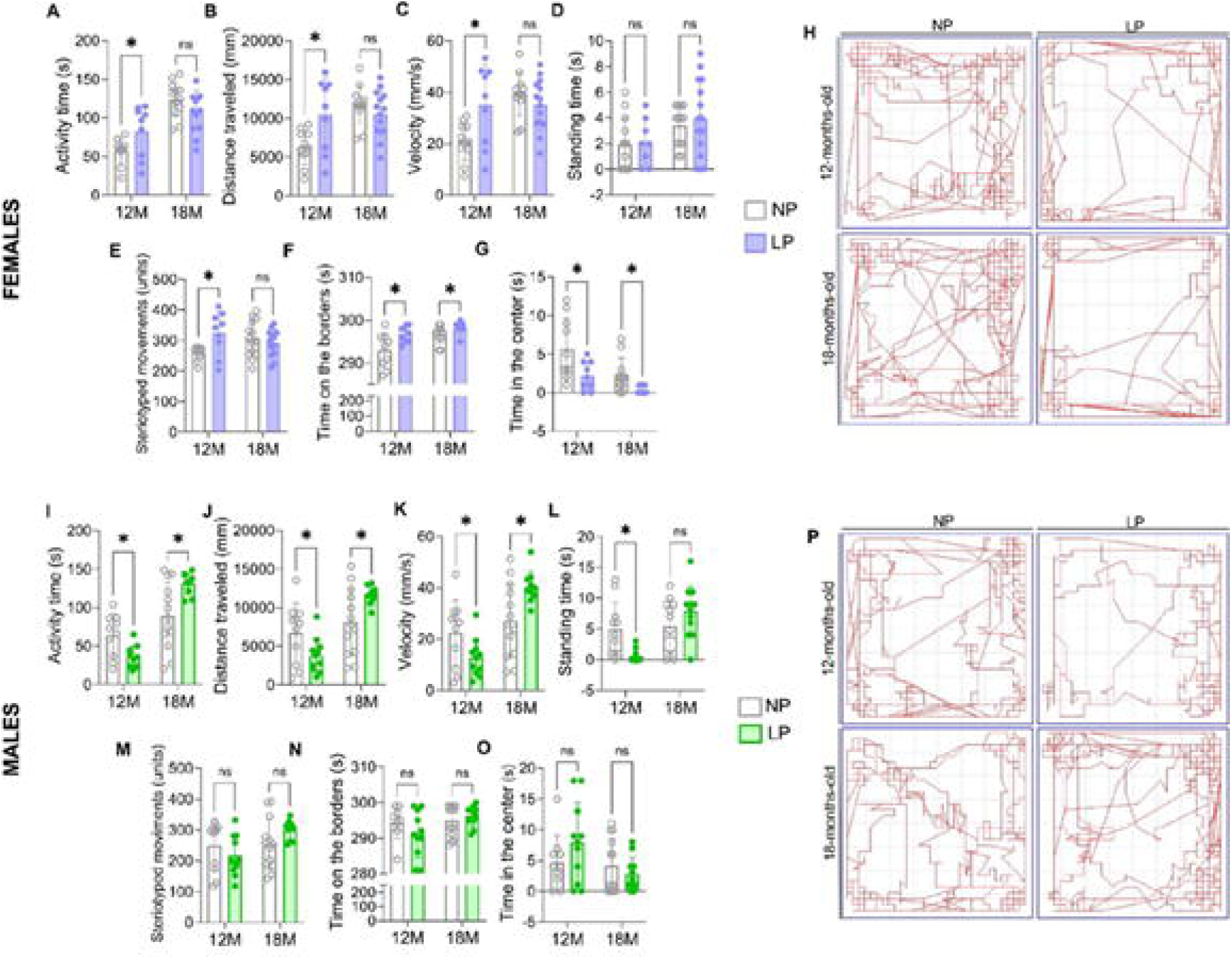
Locomotor and exploratory behavior in the open field (activity monitor) at 12 and 18 months. Females (A–G): (A) activity time, (B) distance travelled, (C) velocity, (D) standing (rearing) time, (E) stereotyped movements, (F) time at the borders and (G) time in the center; (H) representative tracks. Males (I–O): same parameters; (P) representative tracks. NP, normal-protein; LP, low-protein. Data are mean ± SD. Multiple Student’s t-tests; *p < 0.05; ns, not significant.

In males, the pattern was reversed across ages. At 12 months, LP males showed less activity time, a shorter distance travelled, lower velocity (Fig. 3I–K) and less rearing (Fig. 3L) than NP males, indicating reduced horizontal and vertical exploration (Fig. 3P). At 18 months, LP males showed greater activity time, distance travelled and velocity than NP males (Fig. 3I–K), with no difference in rearing. Stereotyped movements, time at the borders and time in the center did not differ between male groups at either age (Fig. 3M–O) (Supplementary parameters Fig. S2).

### 3.4. Sucrose preference and elevated plus-maze

Sucrose preference was not affected in LP females at 12 months but was significantly reduced at 18 months (Fig. 4A). The opposite age pattern was observed in males: LP males showed significantly lower sucrose preference at 12 months, and no difference was detected at 18 months (Fig. 4E). Anhedonia-like behavior therefore emerged in midlife in males and in old age in females.

**Figure 4.**
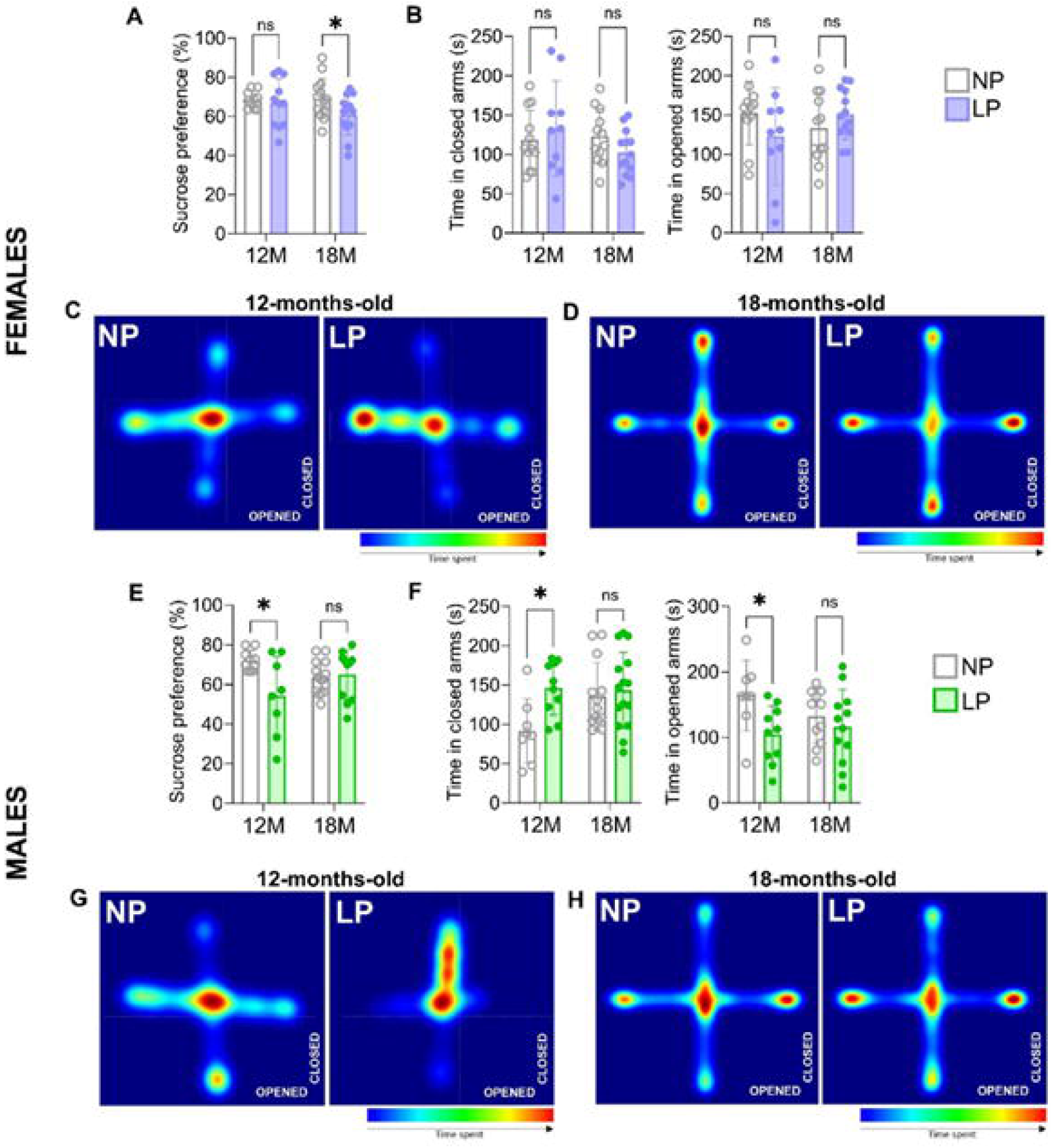
Sucrose preference test (SPT) and elevated plus-maze (EPM). (A, E) Sucrose preference of female and male offspring, respectively. (B, F) Time spent in the closed and open arms of the EPM by females and males; representative heat maps of 12- and 18-month-old females (C, D) and males (G, H). Data are mean ± SD. Multiple Student’s t-tests; *p < 0.05; ns, not significant.

In the EPM, LP females did not differ from NP females in open- or closed-arm time at either age (Fig. 4B–D). At 12 months, LP males spent significantly less time in the open arms and more time in the closed arms than NP males, indicating increased anxiety-like behavior (Fig. 4F, G); no differences were observed at 18 months (Fig. 4F, H). The coexistence of anhedonia-like and anxiety-like behavior with reduced exploration in 12-month-old LP males suggests that their hypolocomotion reflects an affective rather than a motor phenotype. In contrast, 18-month-old LP females showed anhedonia-like behavior without changes in locomotion or EPM indices (Supplementary parameters Fig. S3).

### 3.5. Sociability and social novelty

All groups at both ages spent significantly more time investigating S1 than the empty cup (all *p* < 0.0001), and the sociability index did not differ between NP and LP animals of either sex or age (Fig. 5A, B, E, F, I, J, M, N). Social novelty, which in this protocol indexes short-term social recognition, was affected in females only. At 12 months, NP females unexpectedly spent more time with the familiar S1 than with the novel S2 (*p* < 0.001), whereas LP females showed no preference, resulting in a higher social novelty index in LP females (*p* < 0.01; Fig. 5C, D). At 18 months, NP females no longer discriminated between S1 and S2, consistent with age-related decline in social recognition, whereas LP females preferred S2 (*p* < 0.001), yielding a significantly higher social novelty index (*p* < 0.05; Fig. 5G, H). In males, both groups preferred S2 at 12 months (NP *p* < 0.001; LP *p* < 0.01), and neither group discriminated at 18 months, with no difference in the social novelty index at either age (Fig. 5K, L, O, P).

**Figure 5.**
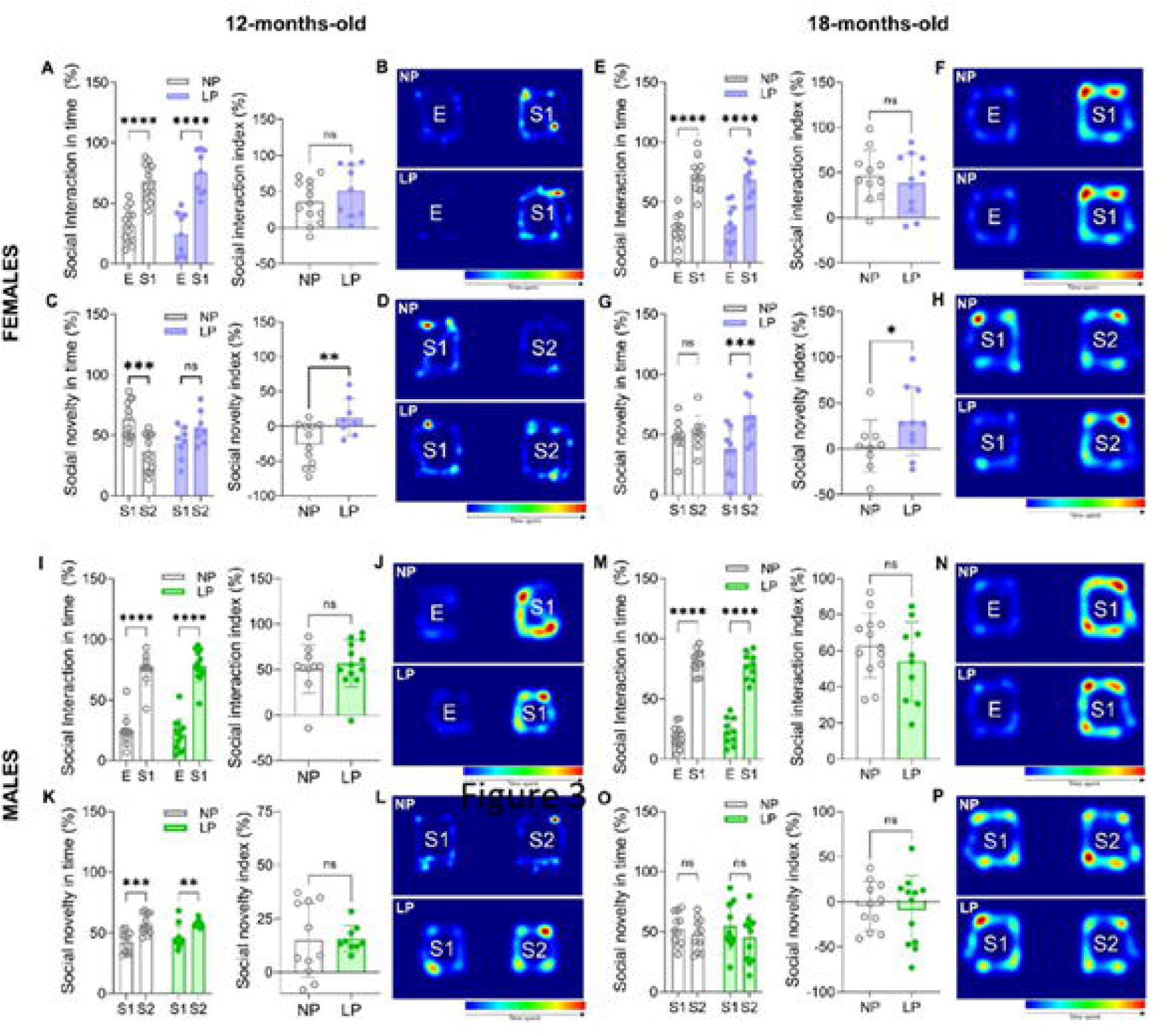
Sociability and social novelty in the three-chamber test. Left, 12-month-old offspring: sociability (A, B, females; I, J, males) and social novelty (C, D, females; K, L, males), with representative heat maps. Right, 18-month-old offspring: sociability (E, F, females; M, N, males) and social novelty (G, H, females; O, P, males). E, empty cup; S1, stranger 1; S2, stranger 2. Data are mean ± SD. Two-way ANOVA (within-group stimulus preference) and Student’s t-test (indices); *p < 0.05, p < 0.01,* **p < 0.001, p < 0.000**

### 3.6. Novel object recognition

#### Females (Fig. 6)

At 12 months, both NP and LP females explored the novel object more than the familiar one in the 3-h test (both *p* < 0.01), without a difference in DI (Fig. 6A, C). In the 24-h test, NP females discriminated the novel object (*p* < 0.05) whereas LP females did not, although the DI did not differ significantly between groups (Fig. 6B, D). At 18 months, neither group discriminated in the 3-h test (Fig. 6E, G). In the 24-h test, NP females failed to discriminate, whereas LP females robustly preferred the novel object (*p* < 0.0001), with a significantly higher DI than NP females (*p* < 0.001; Fig. 6F, H).

**Figure 6.**
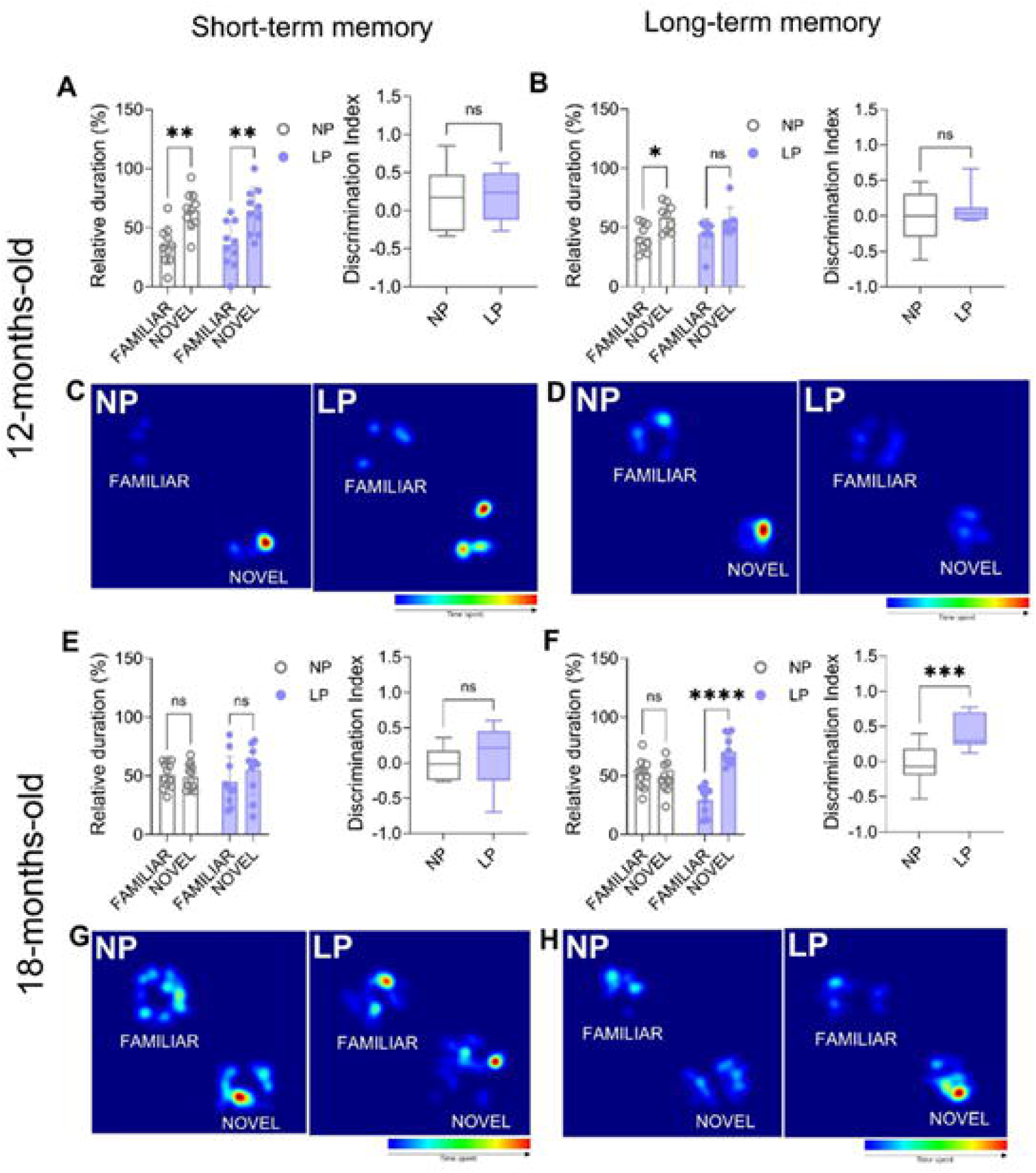
Novel object recognition in female offspring. Relative exploration and discrimination index (DI) in the short-term (3 h; A, E) and long-term (24 h; B, F) tests at 12 (A, B) and 18 months (E, F), with representative heat maps (C, D, G, H). DI is shown as box plots (median, interquartile range, minimum–maximum). Two-way ANOVA and Student’s t-test; *p < 0.05, p **< 0.01,*** **p < 0.001, p < 0.0001; ns, not significan**

#### Males (Fig. 7)

At 12 months, NP males discriminated the novel object in the 3-h test (*p* < 0.05) whereas LP males did not, and the DI was significantly lower in LP males (*p* < 0.05; Fig. 7A, C), indicating impaired short-term recognition memory. In the 24-h test, NP males did not discriminate, whereas LP males preferred the novel object (*p* < 0.05), without a significant DI difference (Fig. 7B, D). At 18 months, neither group discriminated in the 3-h test (Fig. 7E, G). In the 24-h test, NP males spent more time with the familiar object (*p* < 0.05) and LP males showed no preference; the DI was consequently higher in LP males (*p* < 0.05; Fig. 7F, H). This difference reflects a familiarity bias in aged NP males rather than intact recognition in LP males and was therefore not interpreted as a cognitive advantage.

**Figure 7.**
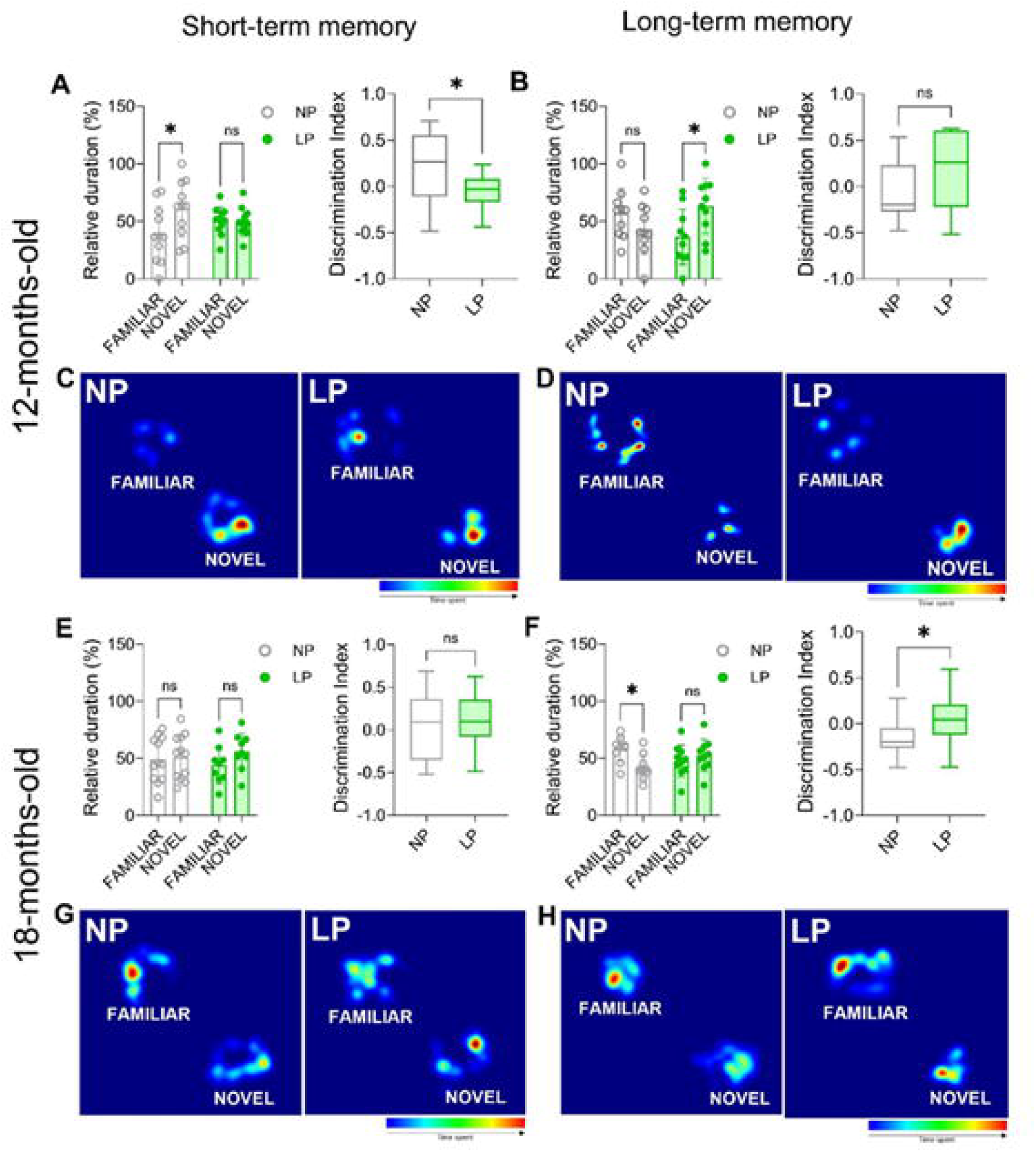
Novel object recognition in male offspring. Relative exploration and discrimination index (DI) in the short-term (3 h; A, E) and long-term (24 h; B, F) tests at 12 (A, B) and 18 months (E, F), with representative heat maps (C, D, G, H). Two-way ANOVA and Student’s t-test; *p < 0.05; ns, not significant.

### 3.7. Spatial learning, memory and cognitive flexibility in the Barnes maze

#### Females (Fig. 8)

Escape latencies decreased across acquisition days in both groups without group differences (Fig. 8A). In the probe trial, LP females spent significantly more time in the target quadrant than NP females (36.5 ± 12.4% vs 26.7 ± 9.4%; *p* = 0.03; Fig. 8A, C), indicating better retention of the escape location. Cognitive scores (Fig. 8B) and the distribution of search strategies (Fig. 8D) did not differ between groups during acquisition. During reversal learning, latencies, probe performance (Fig. 8E, G) and cognitive scores (Fig. 8F, H) were similar between groups.

**Figure 8.**
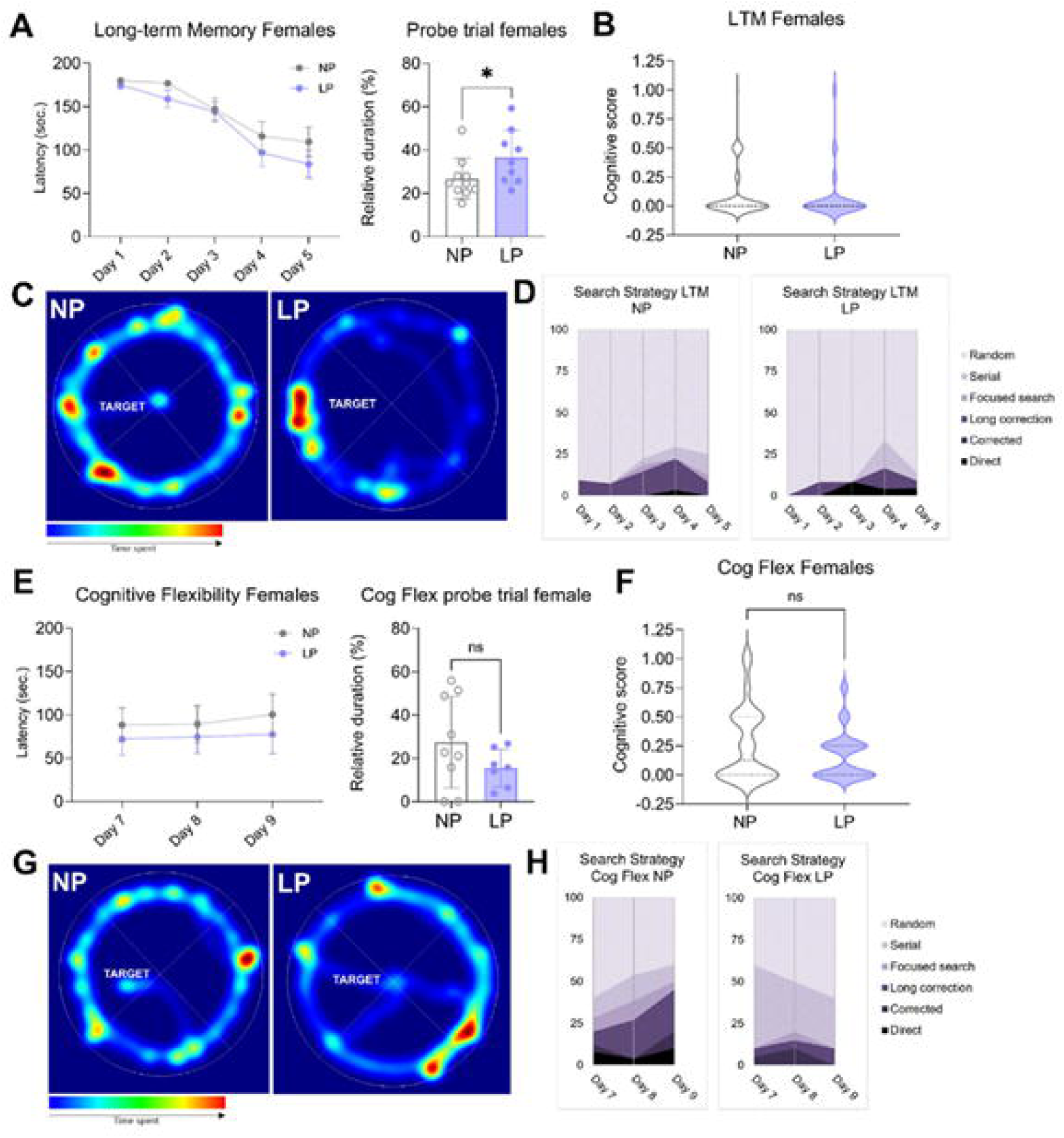
Barnes maze performance of 18-month-old female offspring. (A) Escape latency across acquisition days and target-quadrant time in the probe trial; (B) cognitive score (search-strategy efficiency) during acquisition; (C) representative probe-trial heat maps; (D) daily distribution of search strategies during acquisition. (E) Escape latency during reversal learning and reversal probe performance; (F) cognitive score during reversal; (G) representative reversal-probe heat maps; (H) distribution of search strategies during reversal. Two-way ANOVA for repeated time points, Student’s t-test (parametric) or Mann–Whitney test (non-parametric); *p < 0.05; ns, not significant.

#### Males (Fig. 9)

Escape latencies did not differ on acquisition days 1–4, with a trend towards longer latency in LP males on day 5 (*p* = 0.06; Fig. 9A); probe-trial performance did not differ (Fig. 9A, C). Nevertheless, LP males obtained significantly lower cognitive scores during acquisition (NP 0.23 ± 0.2; LP 0.18 ± 0.2; Mann–Whitney U = 24022; *p* = 0.04; Fig. 9B), reflecting greater reliance on less efficient search strategies (Fig. 9D). During reversal learning, LP males showed significantly longer latency on the last reversal day (*p* < 0.05; Fig. 9E), a trend towards lower cognitive scores (*p* = 0.061; Fig. 9F) and a shift towards less efficient strategies (Fig. 9H), without difference in the reversal probe (Fig. 9E, G).

**Figure 9.**
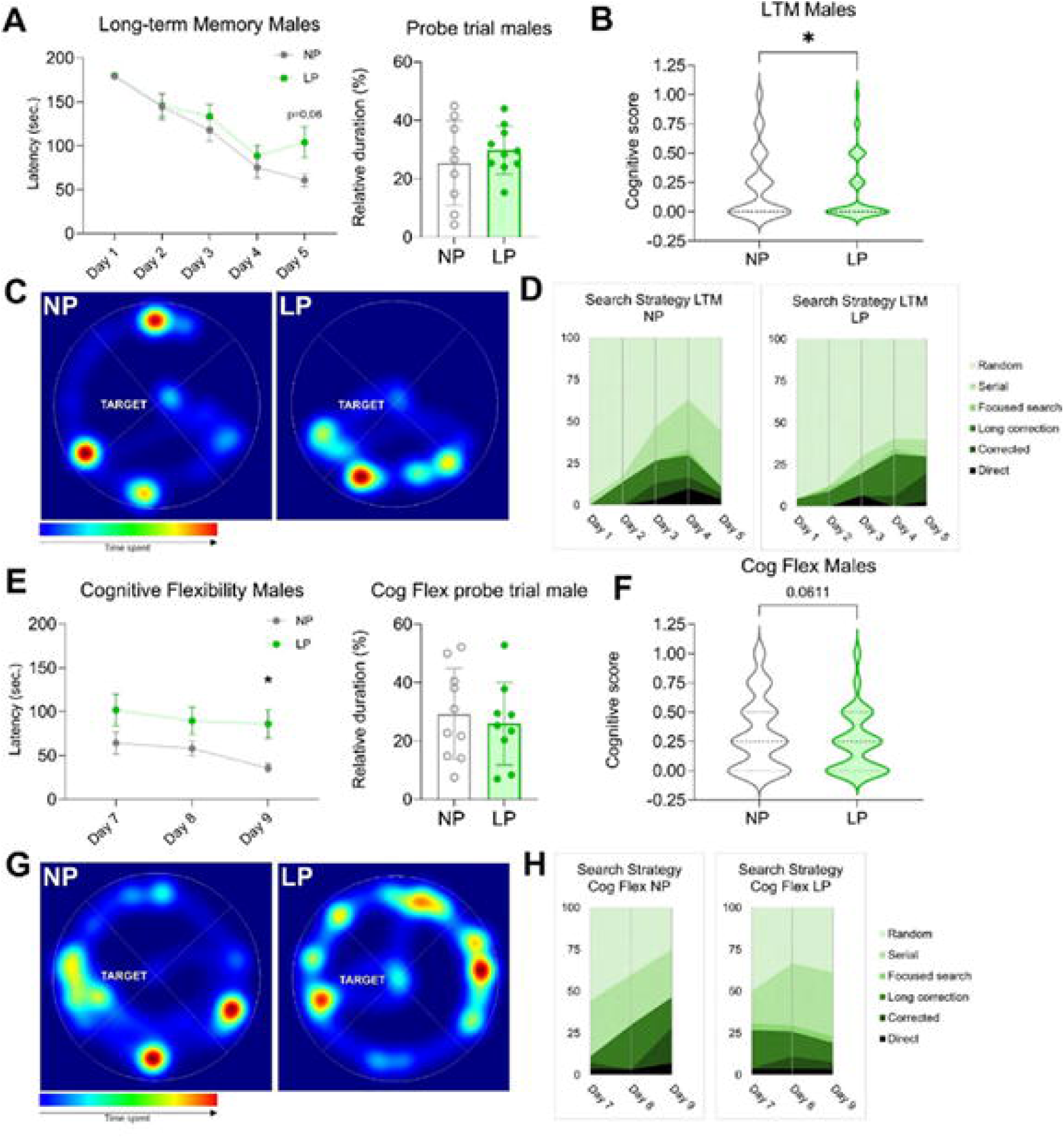
Barnes maze performance of 18-month-old male offspring. Panels as in Fig. 8: (A) acquisition latency and probe trial; (B) acquisition cognitive score; (C) probe heat maps; (D) acquisition search strategies; (E) reversal latency and reversal probe; (F) reversal cognitive score (p = 0.0611); (G) reversal-probe heat maps; (H) reversal search strategies. Two-way ANOVA for repeated time points, Student’s t-test or Mann–Whitney test; *p < 0.05.

### 3.8. Integrated phenotypic map

To visualize the overall pattern, every contrast reported above was coded by direction and functional valence in a single map (Fig. 10A), and within-group stimulus preferences were summarized separately (Fig. 10B). Three features emerge. First, LP effects were domain-specific: sociability was never affected, while each sex showed a distinct set of altered domains. Second, the age of expression differed by sex: alterations in males clustered in affective and short-term mnemonic domains at 12 months and shifted to locomotor and strategic-navigation domains at 18 months, whereas in females anhedonia-like behavior and all three long-term memory advantages appeared at 18 months. Third, the late-life memory advantage of LP females was observed in three independent paradigms (object, social and spatial memory) and occurred precisely where aged NP females had lost discrimination (Fig. 10B). Aged NP males also showed a familiarity bias rather than novelty preference in the 24-h NOR test.

**Figure 10.**
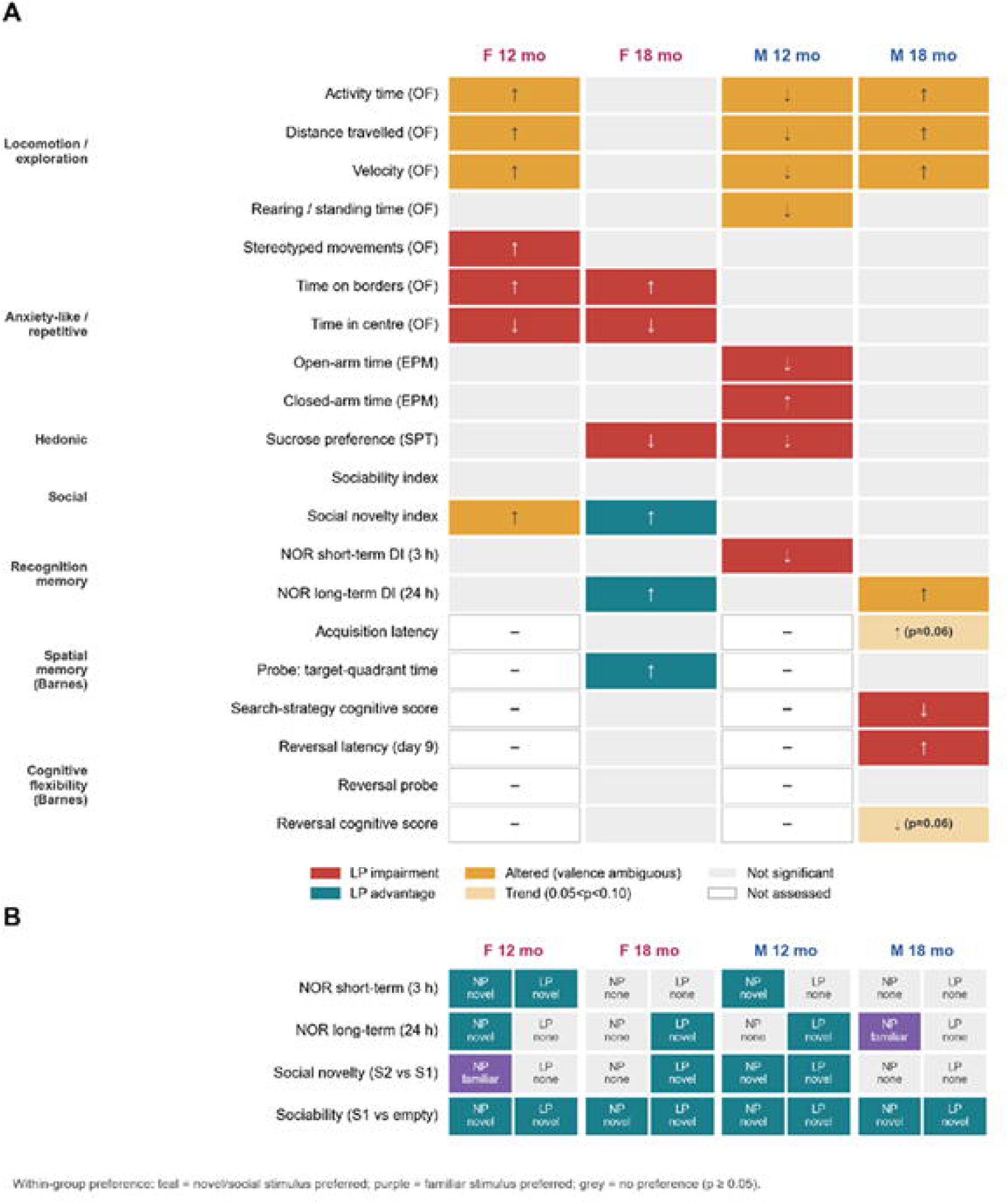
Integrated phenotypic map of the effects of gestational protein restriction on aging offspring. (A) Each cell summarizes the LP-versus-NP contrast reported in Figs. 3–9 for one outcome, sex and age; arrows give the direction of change, and colors give the functional interpretation (red, impairment; teal, advantage; amber, altered with ambiguous valence; light amber, trend 0.05 < p < 0.10; grey, not significant; white, not assessed). (B) Within-group stimulus preference in recognition paradigms (teal, significant preference for the novel or social stimulus; purple, significant preference for the familiar stimulus; grey, no preference). The map is a descriptive synthesis of the reported tests and does not constitute an additional statistical analysis.

## 4. DISCUSSION

This study demonstrates that protein restriction limited to the gestational period programs behavioral trajectories that diverge between the sexes and manifest throughout the second half of life. Males exhibited an early, predominantly affective phenotype (behaviors resembling anhedonia and anxiety, reduced exploration, and impaired short-term recognition at 12 months), which, by 18 months, shifted toward hyperactivity and less efficient spatial search strategies. Females displayed hyperactivity, stereotypies, and thigmotaxis in middle age, and, in old age, behaviors resembling anhedonia; however, they retained long-term memory for objects, social interactions, and spatial information when compared to age-matched control animals. Given that the maternal environment was also altered, these results should be interpreted, more precisely, as the product of a prenatal nutritional insult combined with a modified early postnatal environment. Below, we discuss each of these components, compare our findings with previous studies on protein restriction—as well as with a cohort subjected to prenatal stress, evaluated using the same battery of tests (Fig. 11)—and propose an integrative, testable model (Fig. 12).

**Figure 11.**
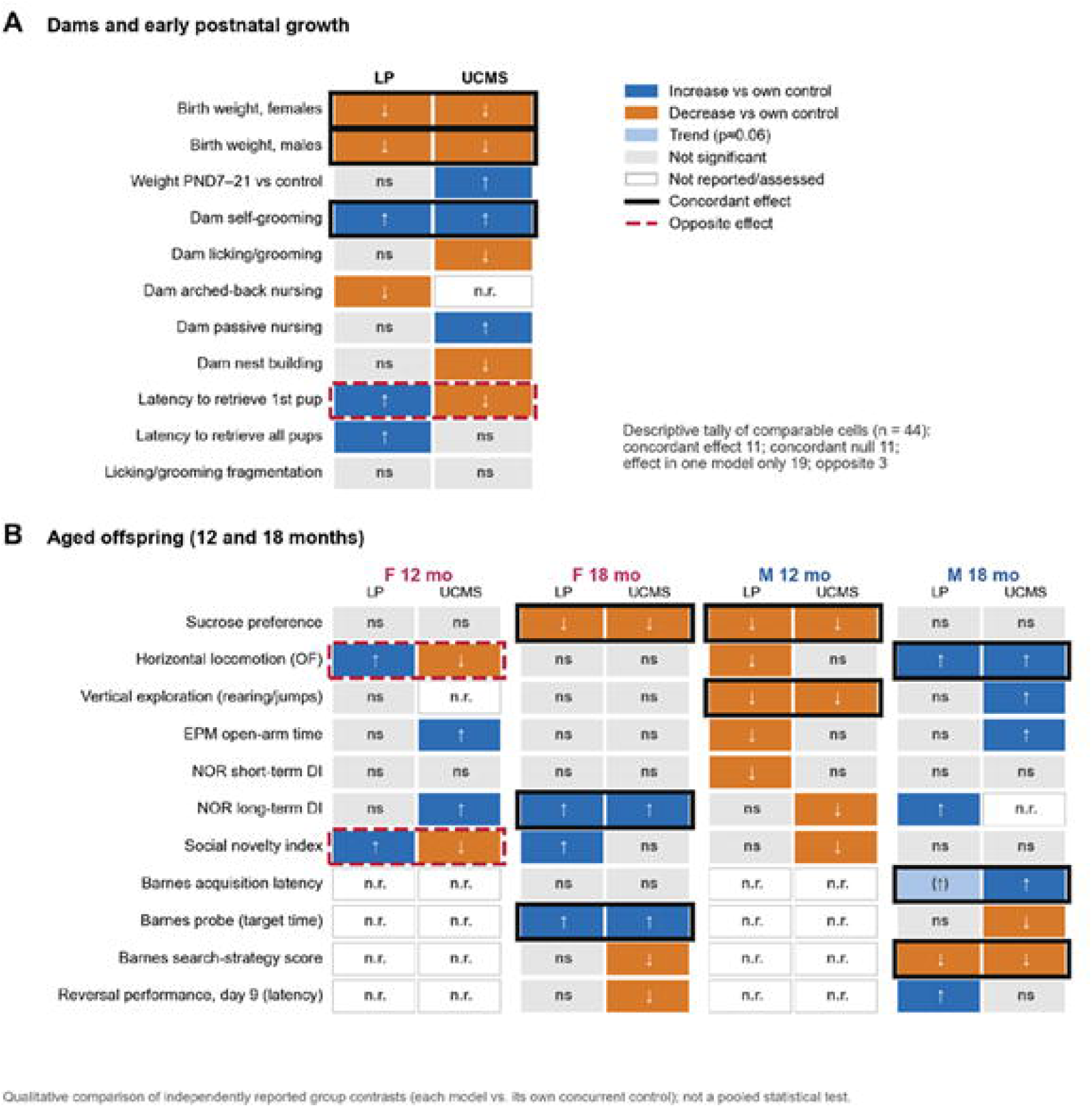
Qualitative cross-model comparison between gestational protein restriction (LP; present study) and prenatal unpredictable chronic mild stress (UCMS; de Souza et al., submitted), both studied in C57BL/6J mice with the same battery and ages in the same laboratory. Each cell pair shows the direction of the treatment effect relative to the respective concurrent control (blue, increase; orange, decrease; light blue, trend; grey, not significant; white, not reported or not assessed). Black frames mark concordant effects and dashed red frames opposite effects. (A) Dams and early postnatal growth. (B) Offspring at 12 and 18 months. The tally (44 comparable cells) is descriptive; no pooled statistical test was performed.

**Figure 12.**
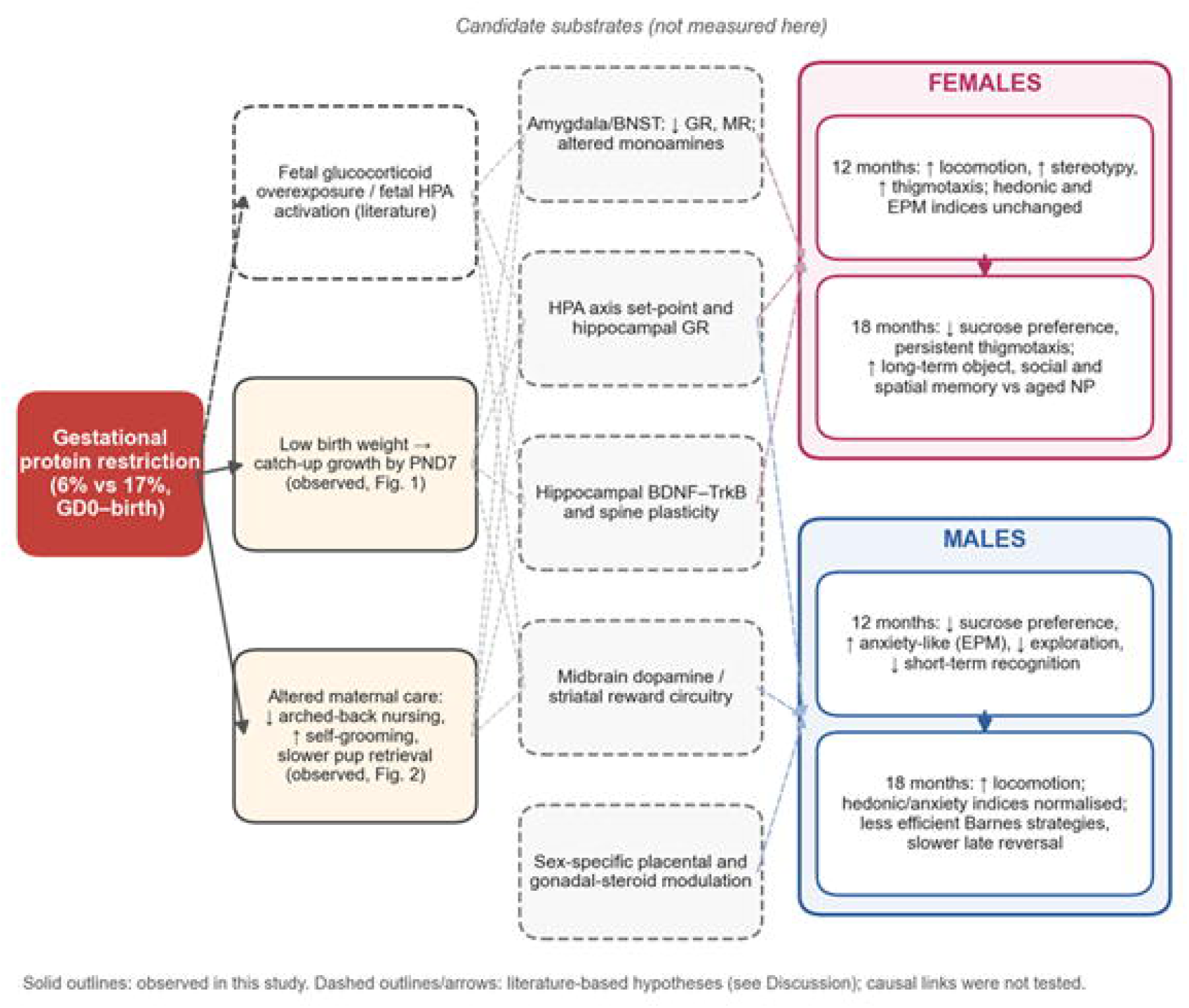
Integrative, hypothesis-generating model. Gestational protein restriction produced intrauterine growth restriction with rapid catch-up growth and altered maternal care (solid outlines; observed). Literature-derived candidate mediators (dashed outlines) include fetal glucocorticoid overexposure (Cottrell et al., 2012), programming of amygdala/BNST and HPA-axis glucocorticoid signaling (Mariano et al., 2025; Torres et al., 2023, 2024), hippocampal BDNF–TrkB signaling (Gutiérrez et al., 2025), midbrain dopaminergic circuitry (Prodani et al., 2024) and sex-specific placental and gonadal-steroid modulation (Bale, 2016; Frick, 2009). These converge on sexually divergent behavioral trajectories from middle to old age (right). Causal links were not tested in this study.

### 4.1. Model validity: growth restriction followed by rapid recovery

The marked reduction in birth weight, followed by complete recovery (“catch-up”) within the first postnatal week (Fig. 1), replicates the core feature of our gestational protein restriction model, previously described in rats by our group (Lamana et al., 2021; Mariano et al., 2025; Sene et al., 2021). The catch-up growth observed here is, in itself, associated with a higher metabolic risk later in life (Hales & Barker, 1992). This contrasts with the persistently lower body weight observed when protein restriction extends throughout the lactation period (Ferroni et al., 2022). Therefore, the timing of the adverse event determines whether the offspring will exhibit a transient or a persistent deficit—a distinction that may account for several discrepancies discussed below.

### 4.2. Alterations in maternal care as a “second hit”

Licking/grooming and its fragmentation, as well as arched-back and passive nursing, were preserved in low-protein (LP) dams; however, these females spent more time self-grooming, in longer bouts, and were slower to retrieve their pups (Fig. 2). Gianatiempo et al. (2020) observed reduced and fragmented maternal care in female mice fed a low-protein diet during gestation and lactation. Using the same model, Alberca et al. (2024) noted maternal behaviors suggestive of anxiety and anhedonia, alongside reduced glucocorticoid receptor (GR) expression in the hippocampus and amygdala. Our data extend these findings, demonstrating that protein restriction during gestation alone is sufficient to alter maternal care. Two non-mutually exclusive mechanisms could explain this persistence: a lasting effect of the gestational deficit on maternal physiology, or effects related to the offspring themselves, as smaller pups may elicit different maternal responses. In contrast, no alterations in maternal care were detected in Wistar rats subjected to gestational protein restriction (Mariano et al., 2025). Interspecies differences in maternal care, as well as variations in observation protocols, may account for this divergence.

The specific profile observed is informative. Increased self-grooming is a sensitive indicator of stress-related, self-directed behavior in rodents (Kalueff et al., 2016). In LP dams, self-grooming was increased both in total duration and in bout length, suggesting a shift in the allocation of maternal time from pup-directed to self-directed activity; the delayed retrieval of displaced pups is consistent with this interpretation. In contrast, arched-back nursing and licking/grooming, the components classically implicated in the maturation of hippocampal GR expression and HPA-axis negative feedback in the offspring (Buschdorf & Meaney, 2016; Caldji et al., 2011), were preserved, making this canonical pathway a less likely mediator of the offspring phenotypes described here. Unlike the prenatal unpredictable chronic mild stress (UCMS) dams in our parallel study, which exhibited fragmented and less predictable care (de Souza et al., 2026), LP dams showed no detectable fragmentation of licking/grooming. Within the framework proposed by Baram and colleagues (Baram et al., 2012; Molet et al., 2016), protein restriction therefore appears to affect the balance between self- and pup-directed behavior more than the predictability of pup-directed care. However, the number of dams analyzed (8–13 per group) limits the sensitivity of this analysis (minimum detectable effect size d ≈ 1.2–1.5; Supplementary Fig. S2), and the absence of cross-fostering prevents the attribution of offspring phenotypes specifically to either prenatal or postnatal factors.

### 4.3. Males: an early affective phenotype followed by a late strategic decline

At 12 months of age, males in the LP (low-protein) group exhibited a combination of the following: anhedonia-like behavior, anxiety-like behavior in the elevated plus-maze (EPM), reduced horizontal and vertical exploration, and impaired short-term recognition (Figs. 3, 4, 7). This set of alterations is consistent with the increased anxiety in the EPM and reduced vertical exploration (*rearing*) observed in young adult C57BL/6J males subjected to maternal proteinrestriction (Crossland et al., 2017); with the increased anxiety-like behavior and elevated corticosterone levels in adult males in our rat-model version (Mariano et al., 2025); and with the anhedonia-like behavior that emerges following perinatal protein restriction in rats (Gutiérrez et al., 2025). From a mechanistic perspective, findings from our group demonstrate that gestational protein restriction alters the neurochemistry of two structures closely linked to persistent anxiety: the amygdala and the bed nucleus of the stria terminalis (BNST). It reduces the content of glucocorticoid receptors (GR) and mineralocorticoid receptors (MR), as well as monoamine levels, in the amygdala; furthermore, it decreases cell count and the levels of GR, MR, and BDNF in the BNST, while simultaneously increasing CRF1 expression during early postnatal life (Torres et al., 2023, 2024). Reduced BDNF–TrkB signaling in the hippocampus and the loss of mature dendritic spines in the CA1 region are associated with anhedonia induced by protein restriction (Gutiérrez et al., 2025). Furthermore, gestational protein restriction promotes the remodeling of dopaminergic neurons in the midbrain and alters dopamine synthesis in the striatum (Prodani et al., 2024). These substrates were not examined in the present study and are proposed as targets for future investigations (Fig. 12).

The normalization of hedonic and anxiety-related indices at 18 months, coupled with the emergence of hyperlocomotion and less efficient search strategies in the Barnes maze, suggests a transition from an affective phenotype to a frontostriatal phenotype. Efficient spatial navigation relies on interactions between the prefrontal and striatal regions, as well as the hippocampus; the prefrontal cortex and dopaminergic systems are particularly vulnerable to aging and prior exposure to stress (Bäckman et al., 2010; McEwen & Morrison, 2013). The longer latency on the final reversal day and the trend toward lower reversal scores point in the same direction. However, the deficit observed in aged males was mild: retention during the probe trial showed no difference, the disparity in acquisition latency was merely a trend, and the effect on strategy scores was small—likely estimated based on individual trials rather than individual animals (see Limitations). An alternative explanation for the apparent normalization of affective indices is that the aging process itself reduces hedonic and exploratory responses in control groups, thereby diminishing the differences between groups. To distinguish genuine recovery from such “floor effects,” longitudinal designs that track the same individuals over time are required.

### 4.4. Females: Mnemonic preservation in later life at a hedonic cost

The trajectory for females was markedly different. At 12 months, females in the LP group exhibited hyperactivity, a higher incidence of stereotypies, and thigmotaxis, yet they showed no alterations in the elevated plus-maze (EPM) test or in sucrose preference (Figs. 3, 4). The dissociation between thigmotaxis in the open field and behavior in the EPM is consistent with the view that these paradigms assess partially distinct constructs: open-field measures are strongly influenced by exploratory drive and locomotor activation, whereas the EPM reflects an approach-avoidance conflict (Prut & Belzung, 2003; Walf & Frye, 2007). Therefore, persistent thigmotaxis may reflect either an altered exploratory strategy or increased anxiety.

The most notable observation is the convergence of results from three independent paradigms at 18 months: females in the LP group performed better than those in the NP (control) group of the same age in tests of object recognition (after 24 hours), social novelty recognition, and memory retention in the Barnes maze (Figs. 5, 6, 8). In each case, the observed advantage occurred because females in the NP group lost their discrimination capacity, whereas females in the LP group retained it (Fig. 10B). Thus, the most accurate description is the preservation of hippocampus-dependent memory against age-related decline, rather than an enhancement beyond the levels observed in young individuals. This finding diverges from reports of premature aging, impaired spatial memory, and increased oxidative stress and senescence markers in female mice exposed to protein restriction during gestation and lactation (Ferroni et al., 2023), as well as from memory and exploratory behavior deficits observed throughout the lifespan in both sexes under the same perinatal protocol (Ferroni et al., 2022). This divergence suggests that the timing of exposure is a critical factor. Restriction limited to the gestational period—followed by rapid growth recovery and normal postnatal nutrition—may trigger adaptive programming; conversely, restriction that persists throughout lactation results in a long-lasting deficit. Consistent with this interpretation, protein restriction limited to the lactation period reduced anxiety-like behaviors and improved spatial memory in male rats (de Oliveira-Silva et al., 2023), while restriction limited to the gestational period in mice also reduced anxiety-like behaviors (Prodani et al., 2024). Three hypotheses—which this study cannot distinguish between—may explain the preservation observed later in life. First, the theoretical frameworks of “match/mismatch” and “stress-inoculation” propose that moderate adversity early in life may prepare an organism for future challenges and promote resilience (Daskalakis et al., 2013; Gluckman & Hanson, 2004; Santarelli et al., 2017); aging may represent one such challenge. Second, ovarian aging may be a contributing factor. In C57BL/6J mice, estrous cyclicity ceases between approximately 11 and 16 months of age (Felicio et al., 1984; Nelson et al., 1982); therefore, the 18-month time point occurs after reproductive senescence in most of these animals. It is worth noting that estradiol is an important regulator of hippocampal memory in females during aging (Frick, 2009; Frick & Kim, 2018). If early-life nutritional programming altered the timing of ovarian aging or the local synthesis of neuroestrogens, hippocampal function might have been preserved in a distinct manner; however, ovarian reserve and estrous cycle status were not assessed in this study. Thirdly, there is the possibility of survival bias, given that perinatal protein restriction has been associated with a marked increase in mortality (Ferroni et al., 2022). If the frailer animals in the LP group died before 18 months of age, the surviving cohort may have been selected for greater robustness. Selection may already have operated around birth: fewer LP pregnancies yielded litters included in the study (33% vs 60% in NP), and cannibalism was more frequent among LP dams. However, survival between weaning and testing was not systematically recorded in the present study, so this possibility could not be evaluated directly.

The preservation of memory was accompanied, at a later stage, by the emergence of an anhedonia-like phenotype, suggesting a trade-off between the hippocampal system and reward systems. Sex differences in striatal dopaminergic function and its sensitivity to estrogen (Becker, 1999), combined with alterations in reward circuitry described following early protein restriction (Gutiérrez et al., 2025; Prodani et al., 2024), make the mesolimbic system a plausible site for this type of hedonic cost. Depressive-like behaviors and changes in hippocampal gene expression have been reported in young female mice following perinatal protein restriction (Belluscio et al., 2016), indicating that females are not protected in the affective domain.

### 4.5. A shared programming signature between nutritional and psychological prenatal adversities

The test battery, strain, ages, and laboratory were identical to those in our study on prenatal unpredictable chronic mild stress (UCMS) (de Souza et al., 2026), which allowed for a qualitative comparison between the two models (Fig. 11). Of the 44 comparable conditions, 11 showed concordant effects, 11 showed concordant null results, 19 showed an effect in only one of the models, and only 3 exhibited opposite effects. The concordant effects are not randomly distributed. In both models, birth weight was reduced in both sexes, dams showed increased self-grooming behavior, and the emergence of anhedonia-like behavior was observed at 12 months in males and at 18 months in females. Furthermore, aged males in both models became hyperactive and employed less efficient strategies in the Barnes maze, whereas aged females in both models maintained long-term memory—for both objects and spatial information—that was superior to that of their respective control groups. The divergences were equally informative. Postnatal growth showed “catch-up” following protein restriction (LP), but an “overshoot” following UCMS; additionally, the two models produced distinct profiles of maternal care. Middle-aged females exhibited hyperactivity following LP, but hypoactivity following UCMS; the models also differed in their results on the elevated plus-maze (EPM) and in their response to social novelty during middle age. We propose that this overlap reflects a core programming signature resulting from prenatal adversity, in which sex and life stage—rather than the specific type of insult—determine which circuits are vulnerable and when. In males, the window of vulnerability occurs in middle age, with a predominantly affective manifestation; in females, it occurs around the time of reproductive senescence, with a predominantly hedonic manifestation and relative preservation of memory. Fetal overexposure to glucocorticoids—a factor common to both psychological stress and protein restriction (Cottrell et al., 2012)—along with alterations in maternal care, constitute the most parsimonious shared causal factors. Sex differences in placental function represent a potential mechanism for these divergences (Bale, 2016; Bale & Epperson, 2015). However, a systematic review found inconsistent evidence for sex-dependent effects of prenatal malnutrition (Christians et al., 2021), and in mice, dietary restriction affected placental gene expression in both sexes, without formal interactions between sex and treatment (Hercus et al., 2025). This comparison is intended solely for hypothesis generation: the cohorts did not share concurrent controls, the effects were not statistically combined, and the study used for comparison has not yet completed the peer-review process.

### 4.6. Translational relevance

In humans, exposure to famine during the prenatal period has been linked to poorer performance in selective attention in later adulthood, which is interpreted as accelerated cognitive aging (de Rooij et al., 2010). Our data suggest that the consequences of prenatal malnutrition may be domain-specific and sex-dependent, rather than causing global impairment; in males, affective vulnerability may precede cognitive alterations, whereas in females, hedonic changes are observed alongside a relative preservation of memory. If confirmed, these findings would indicate a need for sex-stratified and domain-specific monitoring for individuals born following intrauterine growth restriction, with attention paid not only to memory but also to mood and motivation. Environmental enrichment reverses anxiety-like behaviors and molecular alterations induced by perinatal protein restriction in mice (Berardino et al., 2019), indicating that programmed phenotypes are, at least in part, modifiable.

### 4.7. Limitations

Several limitations should be considered when interpreting these findings. First, statistical sensitivity was limited. Given the reported group sizes, most comparisons would only be capable of reliably detecting large-magnitude effects (minimum detectable d ≈ 1.0–1.4 for offspring and ≈ 1.2–1.5 for dams; see Supplementary Fig. S2). The only effect size that could be calculated from the reported summary statistics—retention in the Barnes maze task for females—yielded a Hedges’ *g* of 0.86 (95% CI: 0.10–1.62). Significant results obtained in underpowered studies tend to overestimate effect sizes and have a lower positive predictive value (Button et al., 2013); furthermore, non-significant results cannot be interpreted as an absence of an effect. Second, although including only one offspring per dam in each group avoids pseudoreplication, it would be important to report the number of litters that contributed to each test, as well as the total number of dams per diet, given that litter effects can account for 30% to 60% of phenotypic variance in rodent developmental models (Jiménez & Zylka, 2021; Lazic & Essioux, 2013). Third, ages and sexes were analyzed using separate tests, comparing two groups at a time; factorial or mixed-effects models—incorporating diet, sex, and age—would be required to formally test the interactions suggested by the observed trajectories. Furthermore, no correction for family-wise error was applied when analyzing multiple outcomes. Fourth, the magnitude of the Mann-Whitney U statistic for the cognitive score in males suggests that individual trials, rather than individual animals, were analyzed; mixed-effects models, which account for the animal as a random factor, would be more appropriate. Fifth, neither the stage of the estrous cycle—which modulates social and exploratory behavior in females (Chari et al., 2020)—nor reproductive senescence was monitored. Sixth, the lack of cross-fostering precludes the differentiation between prenatal and postnatal contributions, and maternal behavior was observed only during the light phase. Seventh, Cohort 2 underwent sequential testing, meaning carry-over effects cannot be ruled out; additionally, the 5-minute familiarization period used in the novel object recognition (NOR) test may have been insufficient for aged C57BL/6J mice, which could potentially explain the loss of discrimination ability observed in the aged animals of the NP group. Eighth, survival between weaning and testing was not systematically recorded; therefore, a contribution of selective mortality to the late-life phenotypes cannot be excluded. Finally, no neuroendocrine, neurochemical, or molecular measurements were obtained, and the mechanisms proposed in Fig. 12 remain purely hypothetical.

### 4.8. Future Directions

Key next steps include cross-fostering studies to dissociate prenatal from postnatal effects; longitudinal, intra-individual assessments analyzed using mixed-effects models; evaluation of estrous cyclicity, ovarian reserve, and circulating/hippocampal estradiol levels in aged females; and quantification of hippocampal glucocorticoid receptor (GR) markers, BDNF-TrkB signaling, and striatal dopamine across both age groups. Integrating these findings with brain proteomic data currently being generated by our group, alongside pre-registered replication studies—powered based on the effect sizes reported here—would substantially strengthen causal inference.

## 5. CONCLUSIONS

Protein restriction limited to the gestational period resulted in low birth weight, followed by rapid compensatory growth, and altered specific components of maternal care. It also programmed behavioral aging trajectories that diverged between the sexes. Males exhibited an affective phenotype in middle age, followed by hyperactivity and less efficient spatial strategies in old age. Females displayed hyperactivity and thigmotaxis in middle age, and in old age, they developed anhedonia-like behavior—though they retained long-term memory for objects, social stimuli, and spatial locations, all of which had declined in the control group. Sociability and cognitive flexibility remained largely preserved, pointing to domain-specific rather than global effects. The substantial overlap with a prenatal stress model tested under identical conditions suggests a shared, sex- and age-dependent signature of prenatal adversity. These findings underscore the necessity of treating sex as a biological variable and tracking programmed phenotypes throughout the lifespan.

## 6. Declaration of generative AI and AI-assisted technologies in the manuscript preparation process

During the preparation of this work, the authors used Claude Opus 5.5 (Anthropic) to search and synthesize the literature, integrate the results, draft the manuscript and generate the synthesis figures (Figs. 10–12, Supplementary Figs. S1–S2) and the graphical abstract from the authors’ data. After using this tool, the authors reviewed and edited the content as needed and take full responsibility for the content of the published article.

## 7. Funding

This research was partially funded by the Coordenação de Aperfeiçoamento de Pessoal de Nível Superior (CAPES), the National Council for Scientific and Technological Development (CNPq) [308043/2022-7], Brazilian National Institute of Science and Technology of DOHaD-INCT [409165/2024-7], São Paulo Research Fundation (FAPESP) [2024/06071-1].

## 8. Author contributions (CRediT)

Mariano VS: data curation, project administration, formal analysis, investigation, methodology, writing (original draft); Souza AC: methodology; Grigoletti-Lima GB: methodology, formal analysis, writing (original draft, review and editing); Gontijo JAR: formal analysis, writing (original draft, review and editing); Boer PA: conceptualization, data curation, formal analysis, funding acquisition, investigation, methodology, project administration, resources, writing (original draft, review and editing).

## 9. Data availability

The data supporting the findings of this study are available from the corresponding author upon reasonable request.

## 10. Declaration of competing interest

The authors declare no competing interests.

## Figure Legends

**Supplementary Figure S1.** Experimental design. C57BL/6J dams received a normal-protein (17%) or low-protein (6%) diet from gestational day 0 to birth. Offspring growth and maternal behavior were assessed during lactation; after weaning, independent cohorts of male and female offspring were tested in a behavioral battery (12 and 18 months) or in the Barnes maze (18 months).

**Supplementary Figure *S2*** *-* Supplementary parameters of OF using IR monitor. Supplementary EPM parameters. Oriented locomotor parameters, frequency of standing time, frequency and time of rest events and jumps in OF apparatus of female (A) and male (B) offspring of both experimental groups and ages. NP – normal protein diet; LP – low-protein diet. Multiple Student’s t test was used for statistical analysis. (*) p<0.05.

**Supplementary Figure S3 -** Supplementary EPM parameters. Locomotor parameters, frequency and time in the arms and center of the EPM apparatus of female (A) and male (B) offspring of both experimental groups and ages. NP – normal protein diet; LP – low-protein diet. Multiple Student’s t test was used for statistical analysis. (*) p<0.05.

**Supplementary Figure S4.** Sensitivity analysis. Minimum standardized effect (Cohen’s d) detectable with 80% power at two-sided α = 0.05 for each comparison, given the group sizes reported (NP/LP). The diamond shows the observed effect size (Hedges’ g with 95% CI) for Barnes probe retention in females, the only effect computable from reported summary statistics (36.5 ± 12.4% vs 26.7 ± 9.4%). Dotted lines mark conventional medium (0.5) and large (0.8) effects.

**Table S1.** Dietic components of both experimental chows of the animals.

| Component | NP diet (%) | LP diet (%) |
|---|---|---|
| Cornstarch | 39.67% | 48.0% |
| Casein (84%) | 20.23% | 7.15% |
| Dextrinizade Starch (90-94%) | 13.05% | 15.9% |
| Sucrose | 10.0% | 12.1% |
| Soybean oil | 7.0% | 7.0% |
| Cellulose microfine (fiber) | 5.0% | 5.0% |
| Mineral mix AIN 93 G | 3.5% | 3.5% |
| Vitamin mix AIN 93 G | 1.0% | 1.0% |
| L-cysteine | 0.3% | 0.1% |
| Choline bitartrate | 0.25% | 0.25% |
| BHT (Butylhydroxitoluol) | 0.0014% | 0.0014% |

**Table S2.**
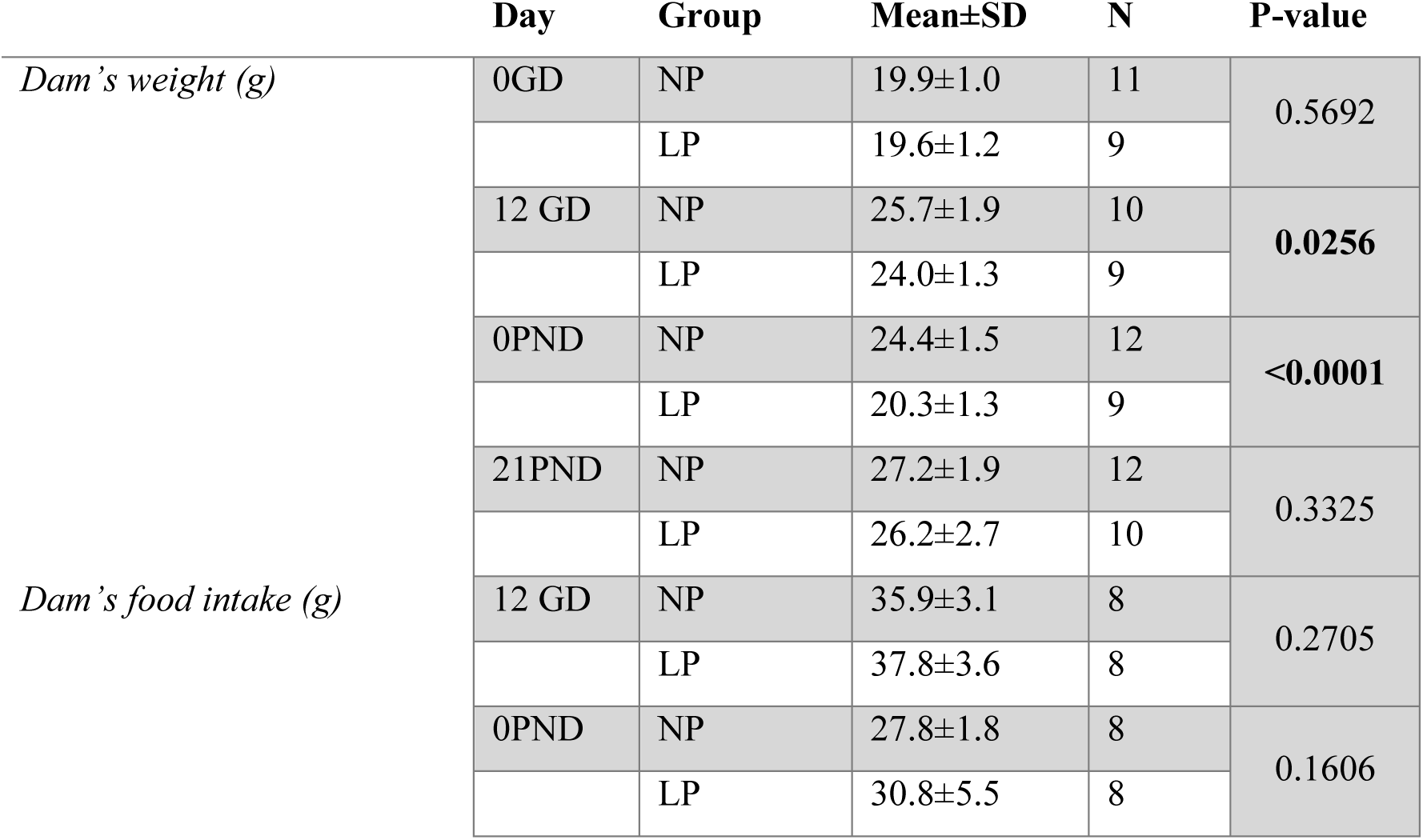
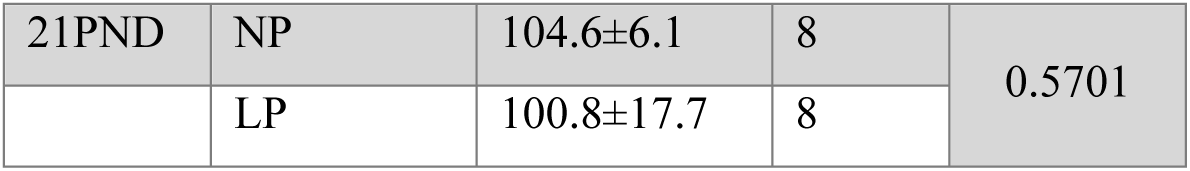
Weekly dam’s weight and intake of both experimental groups.

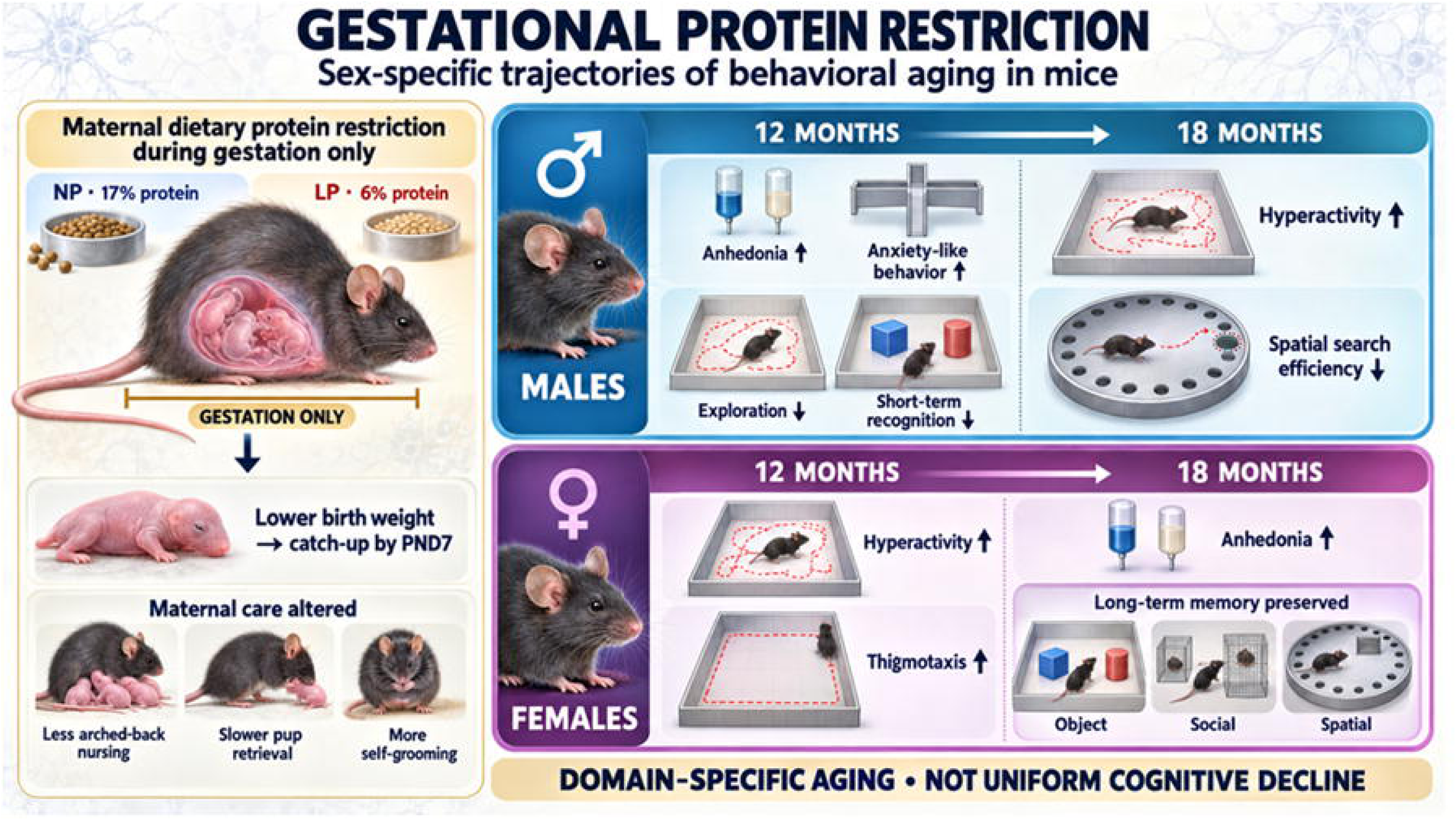

## Notes

### Competing Interest Statement

The authors have declared no competing interest.

