## Supplementary figures and images for "GESTATIONAL PROTEIN RESTRICTION PROGRAMS SEXUALLY DIVERGENT BEHAVIORAL AGING IN MICE: EARLY AFFECTIVE VULNERABILITY IN MALES AND LATE-LIFE MNEMONIC PRESERVATION AT A HEDONIC COST IN FEMALES"

### S1.TIF

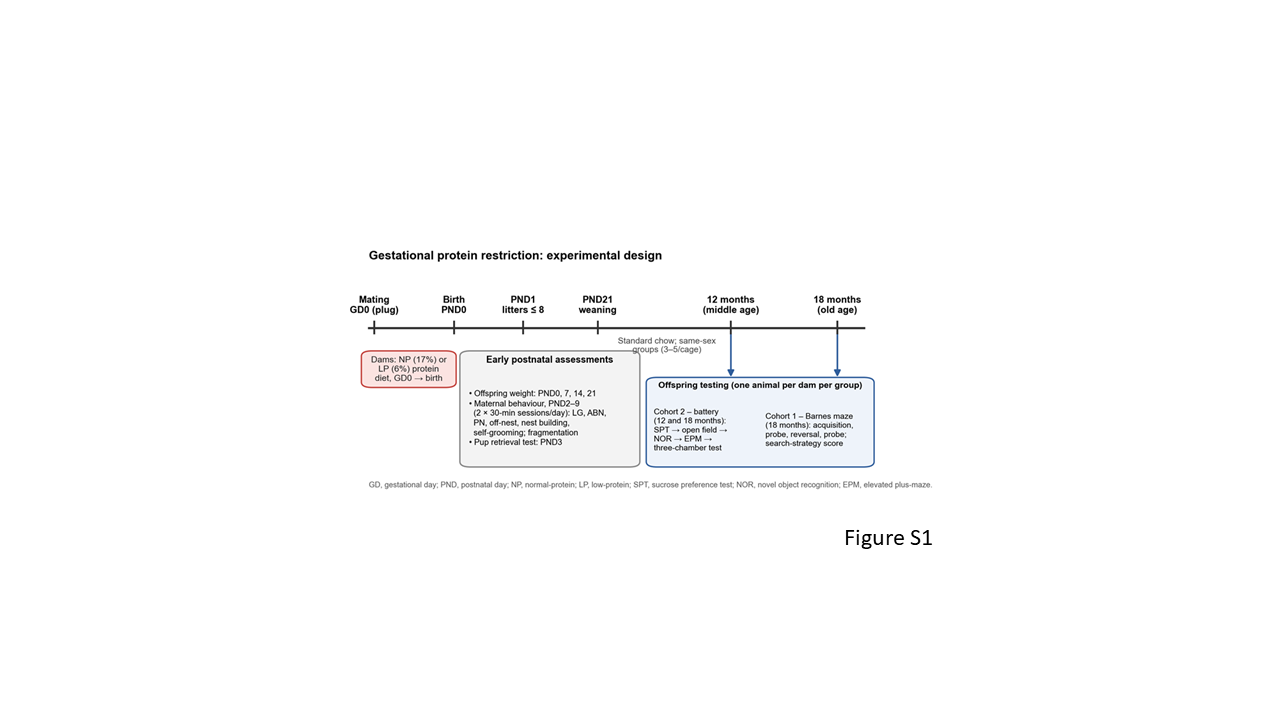

### S2.TIF

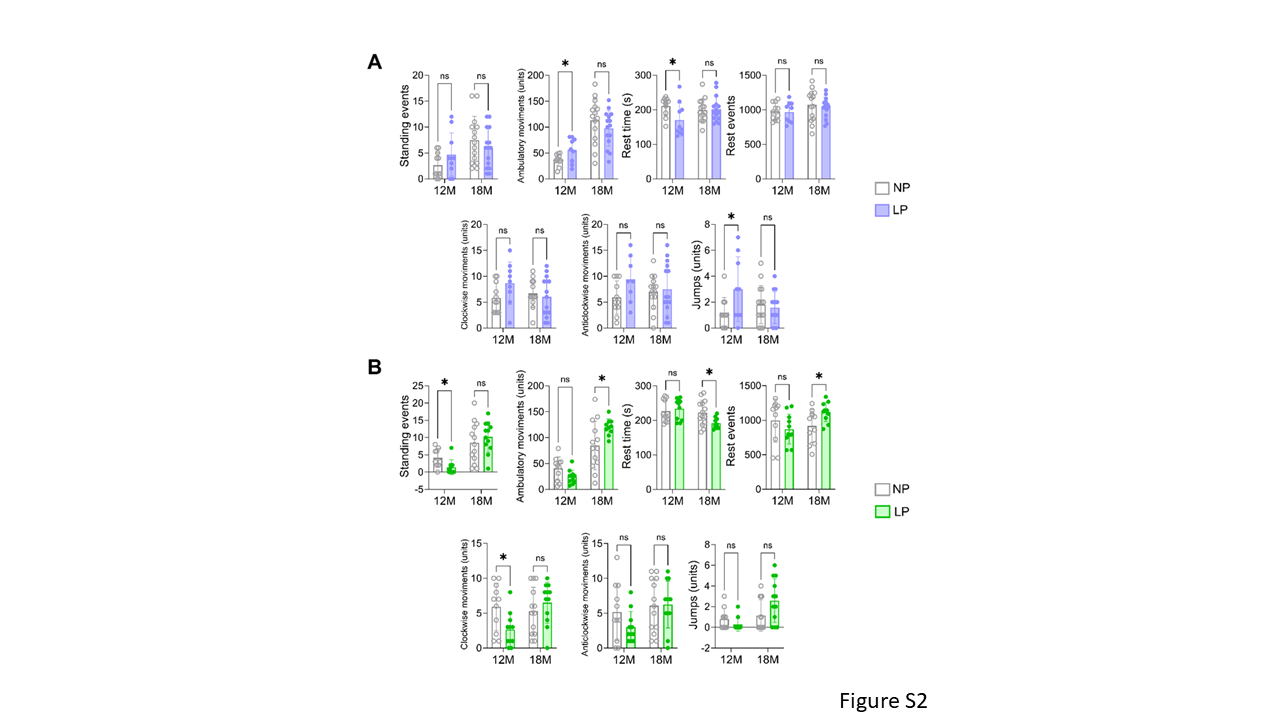

### S3.TIF

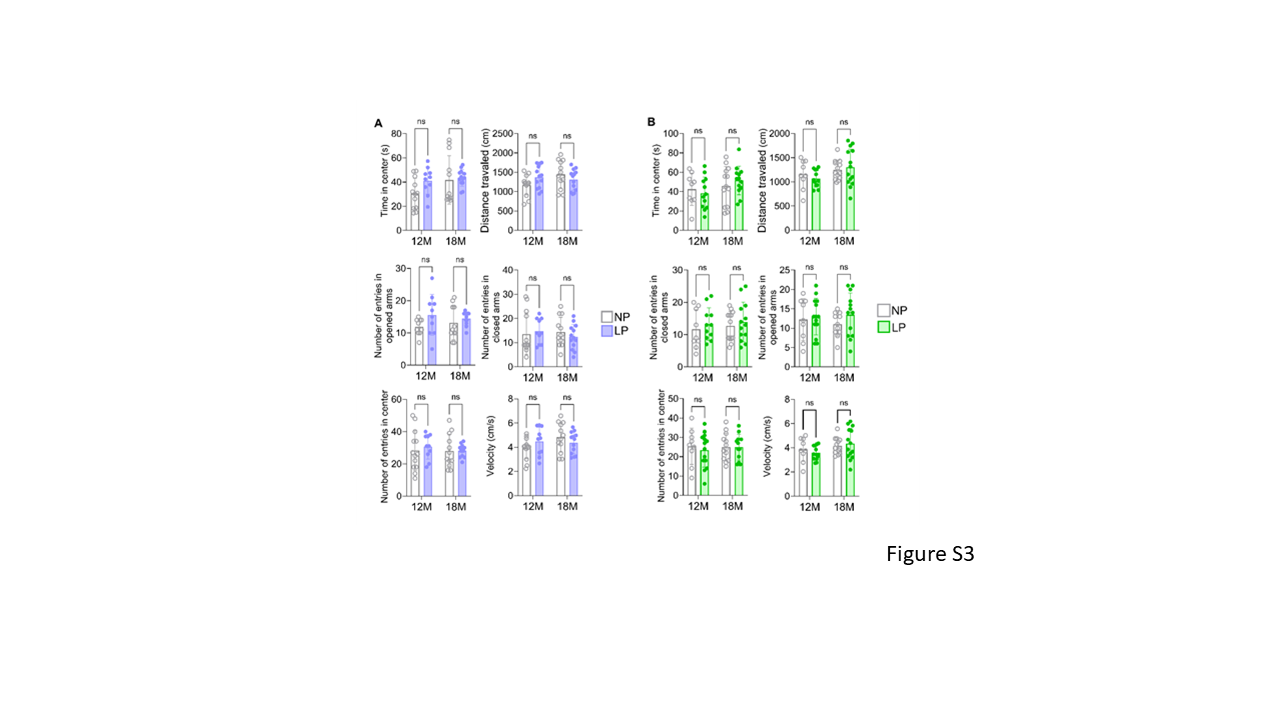

### S4.TIF

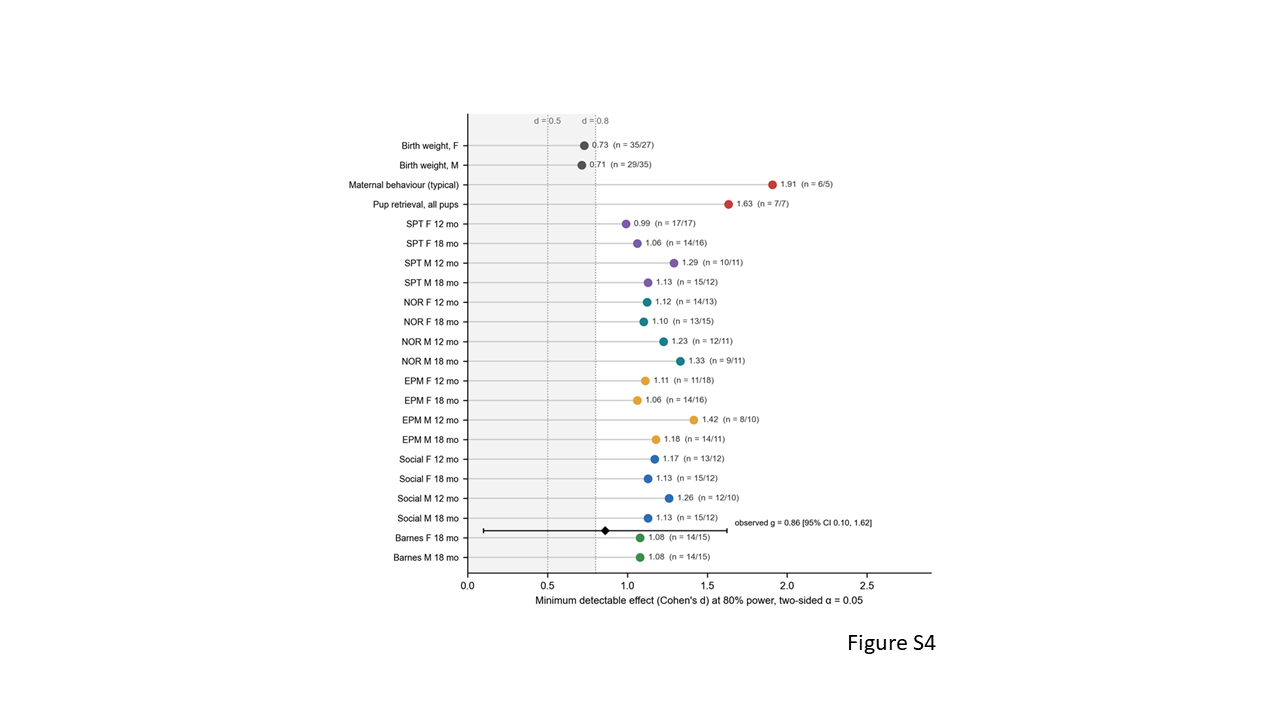
